# Theory of large-bead AFM compression of pressurized rod-shaped bacteria

**DOI:** 10.64898/2026.11.10.765745

**Authors:** Octavio Albarrán, Renata Garcés, Giacomo Po, Christoph F. Schmidt, Jeff D. Eldredge

## Abstract

Motivated by our atomic force microscopy (AFM) experiments measuring turgor pressure in living bacteria, we develop a theoretical description of pressurized spherocylindrical cells compressed by spherical probes whose radius greatly exceeds the cell radius. In this global-loading regime, the mechanical response is governed primarily by cell geometry and turgor pressure, while the elastic properties of the cell envelope enter as higher-order corrections. The strong confinement imposed by the probe drives water efflux from the cytoplasm, thereby increasing the turgor pressure.

We present a theoretical framework for determining bacterial turgor pressure from AFM force– displacement curves, accounting for cell geometry and probe dimensions. Combining numerical simulations with analytical theory, we show that the mechanical response can be approximated in terms of cylindrical and hemispherical contributions. The leading-order force–displacement dependence is captured by an analytical description derived in the large-aspect-ratio limit, where the mechanics reduce to a tension-dominated cross-sectional problem. Finite cell aspect ratio and probe curvature are incorporated through geometric renormalizations. The resulting framework provides a quantitative basis for extracting bacterial turgor pressure from AFM measurements while simultaneously predicting cell deformation geometry.

## I. INTRODUCTION

Bacterial cells maintain internal turgor pressures that can reach several atmospheres, generating substantial mechanical stresses within the cell envelope that play a central role in bacterial shape, growth, and mechanical stability [2, 6, 13, 29, 30, 38]. Quantifying this pressure is essential for understanding the physical state of bacteria and the physiological mechanisms driving its regulation. However, due to the small size of bacterial cells, performing direct measurements in living cells remains experimentally challenging. Atomic force microscopy (AFM) has emerged as a powerful tool for probing bacterial mechanics [9–11, 39, 40], offering high spatial and force resolution. Quantifying turgor pressure from conventional AFM experiments using sharp tips, however, is problematic. It requires solving an inverse mechanical problem in which the localized contact couples geometry, pressure, and nonlinear elasticity [16, 35, 36]. Previous modeling approaches have typically relied on small-deformation perturbation theory and assumed isotropic and axisymmetric geometries [3, 14, 41]. Although recent studies have begun to address nonlinear elasticity and cell-wall anisotropy [1], the effects of these factors for non-axisymmetric geometries under localized indentation remain poorly understood. Moreover, sharp-tip indentation probes a spatially localized region that is difficult to characterize directly through imaging, limiting the available experimental data to force–displacement curves.

In this work, we establish the theoretical and modeling foundation for a technique that will be introduced in a companion publication [12], probing bacterial turgor pressure using large-bead AFM probes to compress rodshaped cells. We focus on the regime in which the probe radius greatly exceeds the cellular radius. For typical material properties of Gram-negative and Gram-positive bacteria, the mechanical response in this global-loading regime is governed primarily by internal pressure and imposed geometry, while the elastic properties of the envelope enter as higher-order corrections. Because the cell envelope is semipermeable and hard to stretch [29, 37], compression drives water efflux from the cell interior, thereby increasing the turgor pressure. The pressure can be inferred from the measured compression force and the contact area between the bacterium and the bead, with the latter determined by modeling. Beyond predicting force–displacement curves and contact areas, the frame-work predicts cell shapes under compression and connects directly to our experimental setup, in which global cell flattening can be captured by high-resolution fluorescence microscopy.

Numerical simulations based on energy minimization of an elastic, pressurized spherocylindrical capsule reveal that the mechanical problem can be simplified by expressing the overall response in terms of cylindrical and hemispherical contributions. The cylindrical contribution, which captures the leading-order force– displacement dependence, is obtained analytically in the asymptotic limit of long capsules and negligible bead curvature. In this case, the system reduces to a crosssectional stress equilibrium, yielding simple relations between force, pressure, and geometry. We then incorporate finite cell aspect ratio and probe curvature through successive geometric renormalizations, preserving the underlying cylindrical relations. The resulting framework provides a unified analytical description of large-bead bacterial compression and establishes a quantitative basis for inferring turgor pressure from AFM measurements.

## II. MECHANICAL FRAMEWORK AND NUMERICAL MODEL

We consider the compression of a single rod-shaped bacterium confined between a rigid substrate and a large spherical bead of radius *R*. The bacterium is modeled as a pressurized sphero-cylindrical capsule of initial length *L* and diameter *D*, with aspect ratio

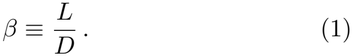

A schematic of the experimental system is shown in Fig. 1, and a three-dimensional representation of the deformed capsule is presented in Fig. 2A. The spherical bead, attached to the AFM cantilever, applies a controlled vertical force *F*, generating global deformation of the capsule.

**FIG. 1:**
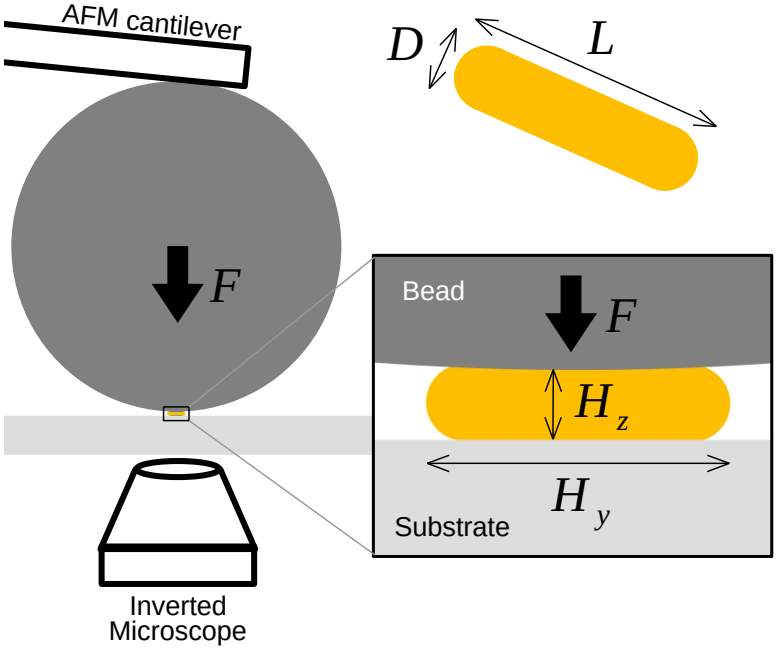
Schematic of the experiment we model. AFM compression of a rod-shaped bacterium modeled as a spherocylinder of length *L* and diameter *D*, indented by a spherical bead of radius *R* applying a force *F*. The schematic preserves the relative proportions *D* : *R* = 1 : 34 used in the numerical simulations in Fig. 2. For experimental details, see [12].

**FIG. 2:**
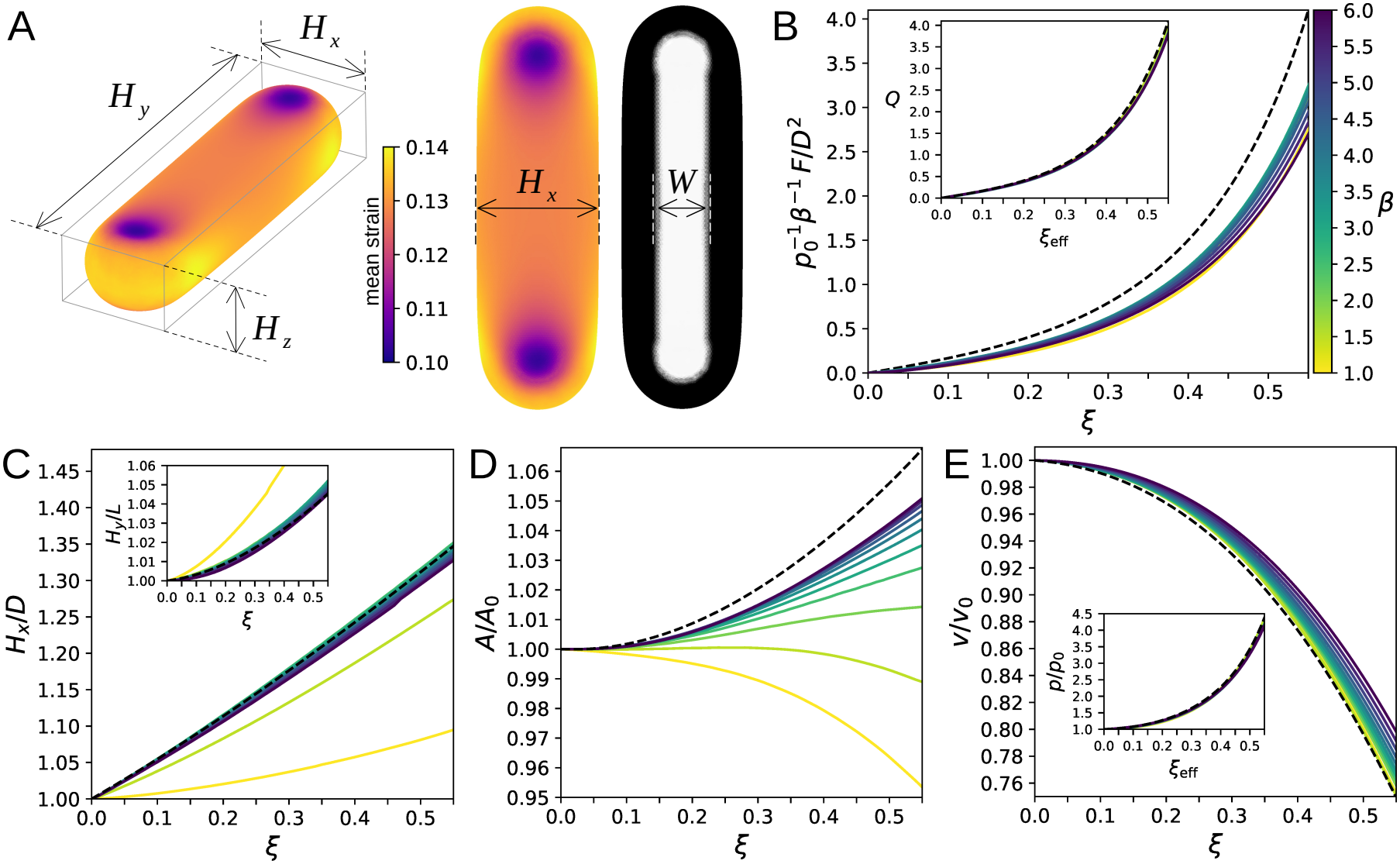
Numerical simulations of spherocylinders compressed by a large spherical bead. Simulations were performed using *Surface Evolver* [5], with a bead radius *R* = 34 *× D* and an elastic model representative of *E. coli* 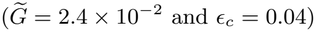. The principal control parameter was fixed at 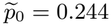, corresponding to a turgor pressure of 1.20 atm for an external osmolality of 200 mOsm*/*L and a pre-strain *ϵ*_0_ = 0.122. **(A)** Compressed configuration of a spherocylinder of aspect ratio *β* = 4.0 and relative compression amplitude *ξ* = 0.30. The local average strain is shown using the *plasma* colormap, while the contact map is shown using a reversed grayscale colormap to indicate the contact force exerted on the capsule. **(B)** Normalized compression force 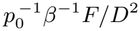 as a function of *ξ* for capsules with different aspect ratios (reverse *viridis* colormap). The dashed line corresponds to the normalized force density 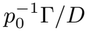 obtained for cylinders in the parallel-plate limit, Eq. (17). Inset: Renormalized force 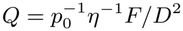, as a function of the effective relative compression amplitude *ξ*_eff_, Eq (39). In panels (B–E), the reverse *viridis* colormap shown in (B) indicates the aspect ratio *β* of the different capsules, ranging from 1.0 to 6.0. Dashed lines denote the corresponding analytical solutions for cylinders in the parallel-plate limit, developed in Sec. IV B. **(C)** Principal dimensions *H*_*x*_*/D* and *H*_*y*_ */D* (inset) as functions of *ξ* (colormap shown in B). **(D)** Relative area *A/A*_0_ as a function of *ξ* (colormap shown in B). **(E)** Relative volume *V/V*_0_ as a function of *ξ* (colormap shown in B). Inset: Relative pressure *p/p*_0_ as a function of the effective relative compression amplitude *ξ*_eff_, Eq (39).

We characterize the deformed shape of the capsule by three geometric measures: the cell height *H*_*z*_ (minimum separation between the bead and substrate), the equatorial longitudinal extension *H*_*y*_, and the equatorial transverse width *H*_*x*_ (see Fig. 2A). In the undeformed state (*F* = 0), these reduce to *H*_*z*_ = *H*_*x*_ = *D* and *H*_*y*_ = *L*.

The cell height *H*_*z*_ is directly obtained from the AFM displacement signal, while *H*_*x*_ and *H*_*y*_ are accessible via optical microscopy. We characterize the compression imposed by the bead through the relative compression amplitude,

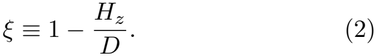

### A. Osmotic Response

The initial volume of the bacterium is

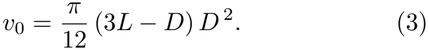

Because the cell envelope is semipermeable and resists area expansion [29, 37], geometric confinement from compression drives passive water efflux from the cytoplasm, reducing cell volume and thereby increasing intracellular osmolyte concentration. As a result, the osmotic pressure difference across the cell envelope, *i*.*e*., the turgor pressure, increases above its initial value. Following the van ‘t Hoff relation [6, 28, 38], the turgor pressure is determined from the cell volume *v* through

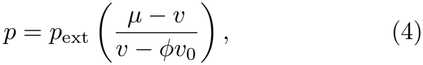

where *p*_ext_ is the ambient osmotic pressure, *µ > v*_0_ is the volume for which the turgor would vanish, and *ϕ* is the fraction of osmotically inactive volume within the cell. Experiments on bacterial osmotic responses support the applicability of this relation. In particular, measurements of the water content of *E. coli* [6, 42] demonstrate the validity of the van ‘t Hoff relation with *ϕ* = 0.37.

To identify the relevant parameters of the system, we formulate the numerical model in dimensionless form. Natural scales are provided by the ambient osmotic pressure *p*_ext_ and the relaxed (unpressurized) cell diameter *D*_rest_. Dimensionless quantities defined using these scales are denoted with an upper tilde. For example, the AFM loading force *F*, cell height *H*_*z*_, volume *v*, and pressure *p* become 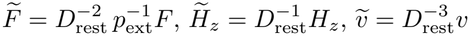, and

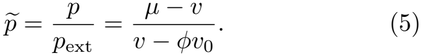

The initial dimensionless turgor,

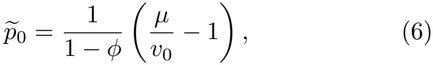

characterizes the initial state of the cell and constitutes the principal control parameter of the system.

Although the theoretical framework is intended to capture generic features of pressurized rod-shaped bacteria, parameter values and modeling assumptions are guided by experimental observations on *E. coli*, which here serves as the principal physiological reference system for both osmotic and mechanical properties.

Because the envelope of the pressurized capsule is already under tension before AFM compression, we introduce an initial pre-strain. For simplicity, we assume that this pre-strain is isotropic, in agreement with previous studies on *E. coli* [22, 24, 25, 28, 29]. With this assumption, the aspect ratio *β* remains independent of pressure, and the isotropic pre-strain is defined as

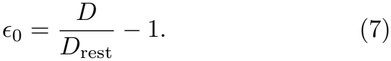

### B. Capsule Simulations

We simulate the mechanics of the pressurized capsule using an energy-minimization framework for elastic surfaces, which is well suited to large-deformation problems with coupled pressure–geometry interactions. The equilibrium configuration is obtained by minimizing the total free energy of the system, which includes osmotic effects, envelope elasticity, and external mechanical loading. We perform the energy minimization using *Surface Evolver* [5].

The total dimensionless free energy of the system is

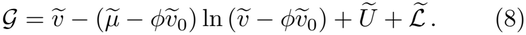

The first two terms represent the osmotic energy of the intracellular fluid, from which the turgor pressure emerges as a thermodynamic response to volume changes, 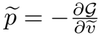. The term *Ũ* represents the elastic energy of the cell envelope. It is constructed from the spring-network model of the *E. coli* cell wall developed in Ref. [1], which accounts for elastic anisotropy and strain hardening (see Appendix A 1). The final term 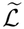 represents the external mechanical work associated with the loading process, with the solid-contact constraints modeled through prescribed contact-force fields.

### C. Parameters and Initial Configuration

The initial geometry and mechanical state of the cell are characterized by the control parameters *β*, 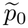, and *ϵ*_0_, corresponding to the aspect ratio, initial dimensionless turgor pressure, and pre-strain, respectively. Other mechanical quantities are determined by these control parameters. For instance, as shown in Appendix B, a variational formulation of the elastic energy *Ũ* yields the following relation for the cell envelope shear modulus:

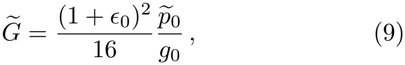

where

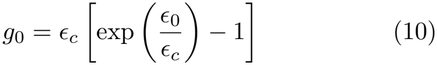

is the initial value of the nonlinear response function describing the elastic behavior of the *E. coli* sacculus (see Appendix A 1). The parameter *ϵ*_*c*_ is the characteristic strain for nonlinear hardening [1]. The dimensionless shear modulus 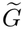 is uniform across the cylindrical body and hemispherical endcaps as a consequence of the assumed isotropic pre-strain.

By varying *ϵ*_*c*_, the exponential response function (10) can be used to describe a broad range of elastic responses, ranging from highly nonlinear elastic cases (*ϵ*_*c*_ ≪ *ϵ*_0_) to the linear limit *ϵ*_*c*_*→ ∞*. In Appendix F, we investigate how the choice of *ϵ*_*c*_, and hence the particular elastic model considered, affects the system. For *E. coli*, the value *ϵ*_*c*_ = 0.04 has been estimated in Ref. [1]. Other representative values for *E. coli* include *D*_rest_ ~ 1.0 *µ*m and *G* ~ 10^−2^ N*/*m [9, 39]. For a representative external osmolality of 200 mOsm*/*L, corresponding to *p*_ext_ ≈ 4.9 atm, these values imply

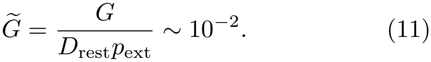

Combining this estimate with Eq. (9) gives 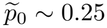 for a physiological pre-strain of approximately 10% [22, 24, 25, 28, 29].

### III. SIMULATION RESULTS

The objective of this section is to identify the principal trends in the cell response through simulations and use these findings to guide the analytical treatment that follows. Unless otherwise stated, we consider a bead radius *R* = 34 *× D* and a characteristic strain for hardening *ϵ*_*c*_ = 0.04. We set the control parameters to 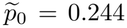, corresponding to an initial turgor pressure of 1.20 atm for an external osmolality of 200 mOsm*/*L, representative of *E. coli*. Simulations are performed over the range of aspect ratios 1 ≤ *β* ≤ 6 (Fig. 2). This range includes the spherical limit (*β* = 1) and encompasses the aspect ratios typically reported for *E. coli* during the cell cycle, 2.5 ≤ *β*≤ 5.5, with an average of *β* = 4.0 [21].

Fig. 2A illustrates a representative compressed state of a capsule with aspect ratio *β* = 4.0 at a compression amplitude *ξ* = 0.30. The strain color map reveals a pronounced contrast between the cylindrical body and the hemispherical endcaps. Along the cylindrical region, the strain field remains nearly uniform. In contrast, the hemispherical caps exhibit strongly heterogeneous strain distributions characteristic of shells with non-zero Gaussian curvature [18, 33, 34, 36], where stretching and bending are coupled. This sharp difference in strain fields motivates an approximation treating the cylindrical body and the hemispherical endcaps separately. The nearly uniform deformation of the cylindrical body makes it possible to describe the local stress equilibrium using a one-dimensional cross-sectional model.

To examine the effect of cell length on the force– displacement curves, we plot the force *F* normalized by the characteristic scale *p*_0_*LD* = *p*_0_ *β D*^2^ for capsules of different aspect ratios (Fig. 2B). This normalization largely eliminates the dependence on length, but a systematic variation remains. As shown in the inset, however, the data collapse onto a single master curve when expressed in terms of the renormalized force *Q* and the effective compression amplitude *ξ*_eff_ (formally defined in Secs. V and VI, respectively), which together account for the contributions of the hemispherical caps and the effects of probe curvature. This collapse indicates that the effects of finite cell aspect ratio and probe curvature can be captured through geometric renormalizations, providing the central motivation for the analytical description developed in the following sections.

Fig. 2C shows how the lengths *H*_*x*_ and *H*_*y*_ increase with compression amplitude. For capsules with sufficiently large aspect ratio (*β* ≳ 2.5), both quantities are well described by the model presented in Sec. IV B and Sec. IV C. In particular, the transverse dimension exhibits an approximately linear scaling, *H*_*x*_*/D* ~ *ξ*, over the explored range of compression amplitudes and increases by ~30% at *ξ* = 0.5. This result reflects the fact that a cylinder can widen when flattened without requiring a change in its cross-sectional perimeter. By contrast, the longitudinal deformation of the capsule is strongly coupled to the axial stretching of the envelope. *H*_*y*_ therefore shows a much smaller increase with compression (Fig. 2C inset), with *H*_*y*_*/L* increasing by only a few percent.

The total surface area of the capsules changes relatively little within the experimentally relevant regime of *ξ* ≲ 0.5, but these small changes exhibit a strong dependence on aspect ratio (Fig. 2D). With increasing compression, spherical capsules and short cylindrical capsules decrease their surface area, whereas longer capsules exhibit a net increase. This difference reflects the distinct Gaussian curvatures of the two geometric regions. The hemispherical endcaps have positive Gaussian curvature, so flattening them requires in-plane deformation [26], leading to a reduction in surface area. By contrast, the cylindrical body has zero Gaussian curvature and can undergo flattening through bending alone, without requiring in-plane deformation [7, 34]. In the simulations, the cylindrical body exhibits an increase in surface area as the pressure increases. For sufficiently large aspect ratios, this contribution becomes dominant.

Capsules of all aspect ratios exhibit a monotonic decrease in volume with increasing compression, reaching volume reductions of approximately 20 % at *ξ* = 0.5 (Fig. 2E). The turgor pressure consequently increases by a factor of approximately 3.5 over the same range. Although the volume curves show a weak dependence on aspect ratio, the pressure curves collapse when plotted against the effective compression amplitude *ξ*_eff_, defined in Eq. (39) to account for probe curvature. Since pressure is directly related to volume through Eq. (4), this collapse indicates that the weak aspect-ratio dependence of the volume curves primarily reflects the geometric effect of the probe rather than intrinsic differences in capsule mechanics.

Taken together, the numerical results reveal a hierarchy of geometric contributions to the deformation of a bacterial cell by large-bead compression, providing the basis for the analytical model developed in the following sections.

## IV. CYLINDRICAL BODY MECHANICS

### A. Equilibrium Equation

Because the AFM bead greatly exceeds the size of the bacteria, we begin by considering the parallel-plate limit, where the curvature of the bead is neglected. Corrections arising from the finite bead radius are introduced in Sec. VI. Within this approximation and for sufficiently large aspect ratios, the deformation becomes dominated by the cylindrical body, and the problem reduces to the mechanics of the cross section of the cylinder (see the schematic in Fig. 3A).

**FIG. 3:**
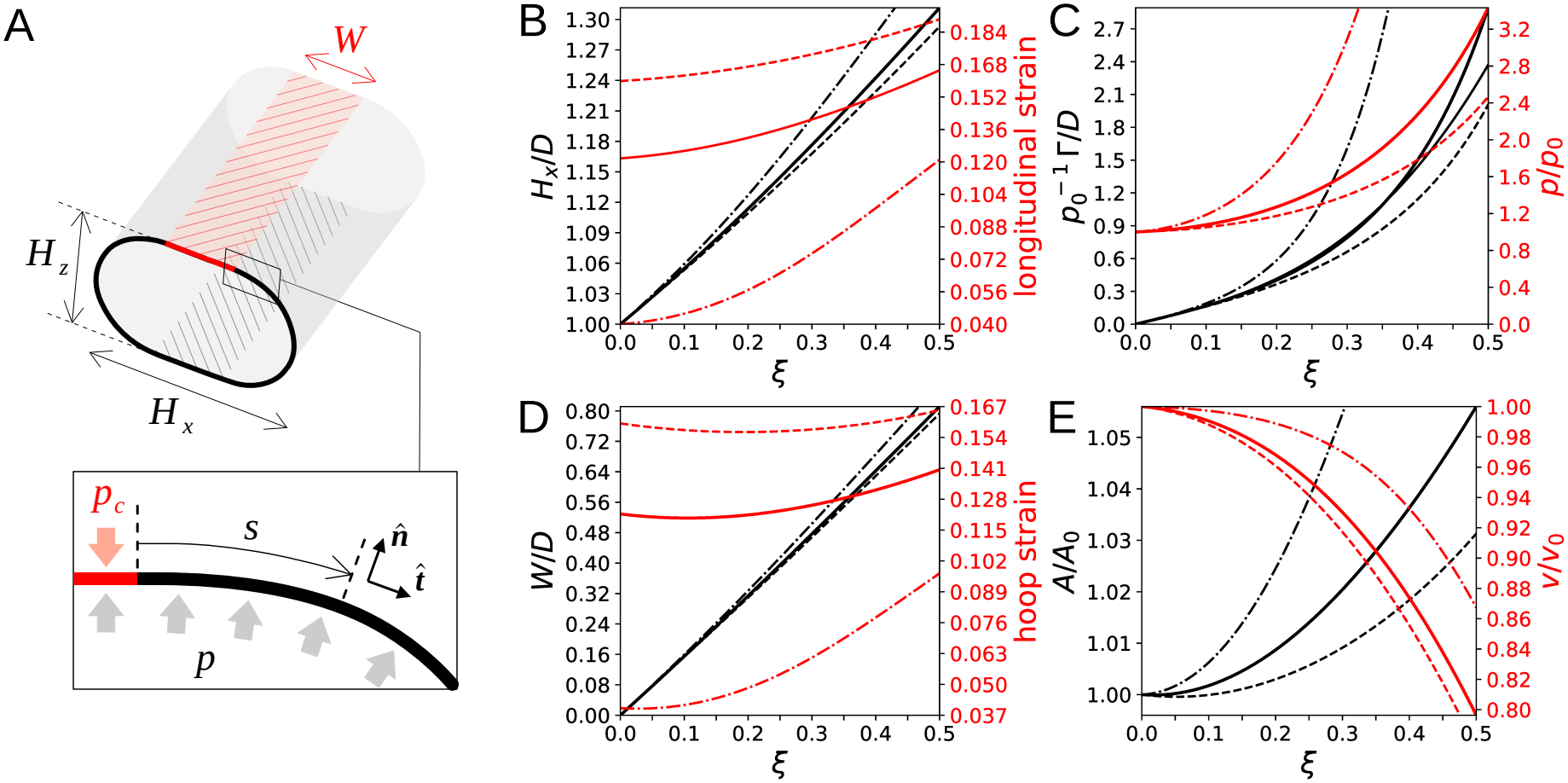
Analytical calculation of parallel-plate compression of the cross section of an infinitely long pressurized cylinder in the tension-dominated limit. **(A)** Schematic of a compressed cylinder. In all panels (B–E), solid lines correspond to 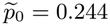 (implying a turgor pressure of 1.2 atm for an external osmolality of 200 mOsm*/*L), dashed lines to 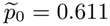 (implying a turgor pressure of 3.0 atm for an external osmolality of 200 mOsm*/*L), and dash-dot lines to 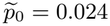 (implying a turgor pressure of 0.12 atm for an external osmolality of 200 mOsm*/*L). All curves are solutions of the system (19) using Eq. (20) and (A12), with the elastic model of Fig. 2 (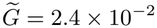and *ϵ*_*c*_ = 0.04). **(B)** Relative transverse width *H*_*x*_*/D* (left y-axis) and longitudinal strain *ϵ*_*y*_ (right *y*-axis) as a function of relative compression amplitude *ξ*. **(C)** Normalized contact force density 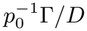 (left y-axis) and relative pressure *p/p*_0_ (right *y*-axis) as a function of relative compression amplitude *ξ*. The gray solid line represents the approximation in Eq. (25) for 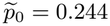.**(D)** Relative contact length *W/D* (left y-axis) and hoop strain *ϵ*_*θ*_ (right *y*-axis) as a function of relative compression amplitude *ξ*. **(E)** Relative surface area *A/A*_0_ (left y-axis) and relative volume *v/v*_0_ (right *y*-axis) as a function of relative compression amplitude *ξ*.

The cross section of the cylinder is modeled as a closed elastic curve characterized by a curvature profile *κ*(*s*), where *s* denotes the arc length. Mechanical equilibrium between the internal forces experienced by the envelope, the pressure *p*, and the load distribution exerted by the parallel plates yields the equilibrium equation (see Appendix C)

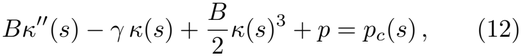

where 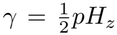 is the hoop stress balancing the force *pH*_*z*_ acting on the lateral free surface and transmitted through the flat contact regions (slip boundary conditions), *B* is the bending stiffness, and *p*_*c*_(*s*) is the contact pressure exerted by the parallel plates. On the free portions of the cross section *p*_*c*_ = 0, whereas in the contact regions *p*_*c*_ = *p*. The latter relation follows directly from the equilibrium equation, since perfect solid contact with the plates imposes *κ* = *κ*^*′′*^ = 0. Furthermore, in the highly pressurized limit, the contact pressure is known to be close to uniform regardless of the shape of the capsule [31, 36]. Consequently, the uniform contact relation, *p*_*c*_ = *p*, can be extended to the full 3D spherocylinder, providing a tractable connection between the loading force and the contact area *A*_*c*_ between the capsule and the bead,

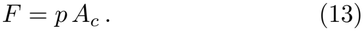

### b. Small-Bending-Stiffness Limit

Over the experimentally relevant range of compressions, *D/*2 *< H*_*z*_ *< D*, the hoop stress scales as *γ* ~ *pD*. In the tension-dominated regime, *γ ≫ B/D*^2^, Eq. (12) predicts that the curvature profile changes rapidly from zero (in the contact regions with the parallel plates) to the constant value 2*/H*_*z*_ in the lateral free portions. Thus, the role of bending stiffness is limited to regularizing this transition by enforcing curvature continuity through a narrow boundary layer near the contact edge. Neglecting the nonlinear term in Eq. (12), one obtains

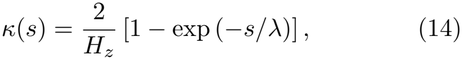

where 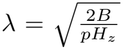 is the characteristic length that separates the flat contact region from the free region of constant curvature. Since *H*_*z*_ ~ *D*, the condition *λ* ≪ *D* that distinguishes the tension-dominated regime becomes

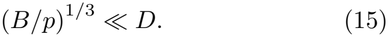

In *E. coli*, the peptidoglycan (PG) *sacculus* is known to consist of a mostly single molecular layer of about 4 nm thickness [15, 32, 37, 39]. Therefore, the bending rigidity of the whole cell envelope is expected to be roughly of the same order of magnitude as that of its lipid bilayers, *B* ~ 10^−19^ N m. Taking *p* ~ 10^5^ N*/*m^2^, we obtain (*B/p*)^1*/*3^~ 10^−8^ m, approximately two orders of magnitude smaller than the bacterial diameter.

In Gram-positive *B. subtilis*, the PG network comprises about ten molecular layers [4, 32]. Estimating the bending stiffness from a Young’s modulus *E* ~ 10^7^ N*/*m^2^ and a thickness *t* ≈ 40 nm [19, 23] yields

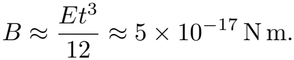

With *p* ~ 10^6^ N*/*m^2^ [30, 38], one again finds (*B/p*)^1*/*3^ ~ 10^−8^ m.

Therefore, for both Gram-negative and Gram-positive rod-shaped bacteria, bending effects remain confined to a negligibly small region near the contact edge. Outside this boundary layer, the cross-section is in a tensiondominated regime, in which the lateral free portions adopt semicircular profiles with diameter *H*_*z*_. The resulting contact length (Fig. 3A) is

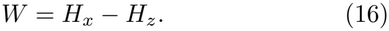

Owing to translational invariance along the cylinder axis, extensive quantities such as force and volume are naturally expressed per unit initial cylinder length. Using Eq. (13), the contact force density Γ is given by

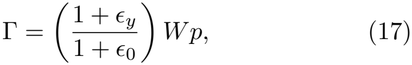

where *ϵ*_*y*_ is the longitudinal strain and *ϵ*_0_ is the isotropic pre-strain (7). Similarly, the cross-sectional area gives the volume per unit current cylinder length. Thus, the volume density *ρ* (volume per unit initial cylinder length) is

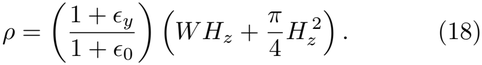

The pressure–volume relation in Eq. (4) can then be expressed in terms of the volume density.

In the tension-dominated limit, the hoop stress *σ*_*θ*_ = *γ* and the longitudinal stress *σ*_*y*_ are uniform and determined by the geometrical relations

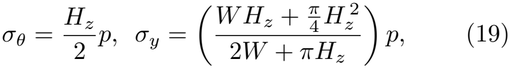

where the quantity in parentheses in the expression for *σ*_*y*_, obtained from a longitudinal balance of forces, corresponds to the enclosed cross-sectional area divided by its contour length.

Because the stress field is uniform in the cross section, the corresponding strain is uniform as well. The hoop strain can therefore be obtained directly from the crosssectional perimeter,

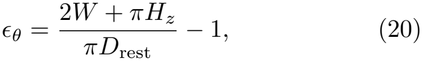

where *D*_rest_ is the resting diameter introduced in (7). Combining Eqs. (20), (19), and the stress–strain relationships (A12) yields a closed algebraic system for *ϵ*_*θ*_ and *ϵ*_*y*_, which can be solved numerically. Fig. 3 illustrates solutions for three representative values of the initial dimensionless turgor pressure, 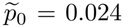, 0.244, and 0.611, corresponding to turgor pressures of 0.12 atm, 1.20 atm, and 3.00 atm, respectively, for an external osmolality of 200 mOsm*/*L.

Fig. 3B,D show the cross-sectional size *H*_*x*_*/D* and the contact length *W/D* as functions of *ξ*, both functions being approximately linear. The right *y*-axes additionally show the principal strains (*ϵ*_*y*_, *ϵ*_*θ*_). Larger values of 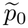 correspond to larger initial strains, *ϵ*_*y*_ = *ϵ*_*θ*_ = *ϵ*_0_.

### C. Flattening Kinematics

The nearly linear response of *H*_*x*_ and *W* to compression is due to the small variation of the cross-sectional perimeter. Rearranging Eq. (20) gives

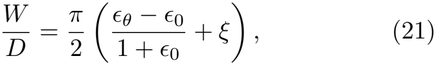

so that when *ϵ*_*θ*_ ≈ *ϵ*_0_,

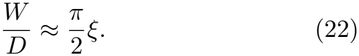

This approximation is largely independent of the constitutive model assumed for the cell envelope and becomes increasingly accurate at larger values of *p*_0_, where the pre-stressed envelope undergoes smaller strain variations. We refer to this geometric behavior as the flattening kinematics of the cylindrical body.

Fig. 2C compares the predictions of the cylindrical model (dashed lines) with the numerical simulations of the complete capsules for *H*_*x*_*/D* and *H*_*y*_*/L*. Good agreement is obtained for *β ≥* 2.0, indicating that, at sufficiently large aspect ratios, the reduced cross-sectional description accurately captures the deformation of the cylindrical body.

The flattening kinematics can be used to derive an approximate contact force density from geometric relations. Assuming *ϵ*_*θ*_ = *ϵ*_*y*_ = *ϵ*_0_, Eq. (18) gives 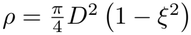.

Consequently, the relative volume is

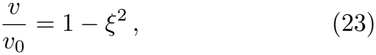

which, through the van’t Hoff relation, gives the relative pressure

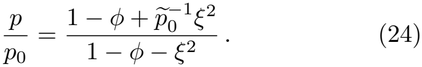

Finally, combining Eqs. (22) and (17) and expanding for small *ξ*, we obtain

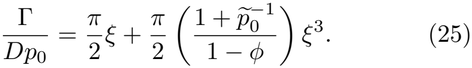

For sufficiently small compression, the force density therefore increases linearly with *ξ*. In this regime, the slope of the loading curve provides a direct measurement of the initial turgor pressure, without requiring knowledge of the mechanical properties of the cell wall. The control parameter 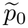 first enters at third order. This correction arises from the pressure increase associated with volume reduction and becomes more significant as 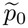 decreases.

In Fig. 3C, the gray solid line depicts Eq. (25) for 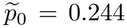, and we see that it approximates the exact solution (black solid line) well up to *ξ* ≈ 0.4, while the linear regime holds for *ξ* ≲ 0.20. To further demonstrate the weak dependence of the flattening kinematics and the linear force response on the constitutive model, Appendix D derives analytical expressions for the strains in the small-compression regime in terms of 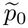 and a single parameter characterizing the elastic response.

It is tempting to use Eq. (25) to determine the turgor pressure directly from the small-compression limit of the experimental curve, *L*^−1^*F* (*ξ*)*/D* = *β*^−1^*F* (*ξ*)*/D*^2^, which is asymptotically equivalent to Γ*/D*. However, the numerical simulations shown in Fig. 2 already indicate that the cylindrical approximation does not fully capture the mechanical response of the capsule. In particular, the force curves in Fig. 2B deviate systematically from 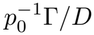 (dashed line), while the corresponding area and volume variations in Fig. 2D, E are likewise not fully described by the cylindrical model alone. As discussed in Secs. V and VI, these deviations arise from both the contribution of the hemispherical caps and the finite curvature of the compressing bead. The following sections address both effects.

## V. DECOMPOSITION OF THE SPHERO-CYLINDRICAL RESPONSE

Next, we extend the parallel-plate compression setup to a spherocylinder of finite aspect ratio *β*. Rather than solving the elastic problem directly, we introduce an approximation based on two asymptotic limits: the spherical solution at *β* = 1 and the cylindrical solution for an infinitely long capsule (*β→ ∞*). The underlying idea is to exploit the natural length-scale decomposition of the capsule to construct a simple geometric interpolation. The cylindrical contributions to force and volume are obtained by multiplying the corresponding densities Γ and *ρ*, defined in Eqs. (17) and (18), by the cylindrical length (*β* − 1)*D*. The spherical solution then provides the remaining contributions to the force and volume. This leads to the approximations

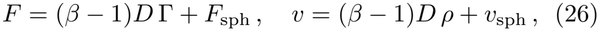

where *F*_sph_ and *v*_sph_ are the force response and volume of the spherical capsule, respectively. By construction, these expressions recover the spherical solution at *β* = 1 and approach the cylindrical limit as the body length increases. Comparison with numerical simulations shows that the approximation reproduces the spherocylindrical response over a broad range of parameters (Appendix G).

### A. Force Master Curve

The force decomposition in Eq. (26) allows the force response to be recast in the master form

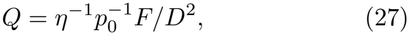

where *η* is the renormalization factor

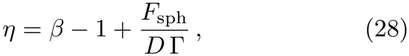

such that *Q* is independent of *β* and equals the normalized force in the cylindrical limit,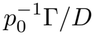.

The quantity *F*_sph_*/*(*D* Γ) in Eq. (28) can be interpreted as the length of an equivalent cylindrical body, normalized by *D*, that generates the same compression force as the two hemispherical caps. Adding this length to the relative length of the actual cylindrical body, *β* − 1, gives the equivalent aspect ratio *η*, which rescales the spherocylinder into an equivalent cylinder and provides a more natural normalization factor than the bare geometric ratio *β*.

Constructing the master curve *Q* requires first solving the compression problem for a spherical capsule. However, obtaining such a solution is challenging, as it involves nonlinear membrane equilibrium equations in which large geometric rotations and strong coupling between meridional and hoop stresses generally preclude a closed-form solution [8, 17, 20, 27]. To our knowledge, no analytical treatment is available for the specific setup considered here, which combines a hyperelastic 2D spring network with internal pressure–volume coupling (4). We therefore use our numerical simulation framework to compute the force *F*_sph_ and volume *v*_sph_ of the compressed spherical capsule, needed to evaluate the model (26).

### B. Numerical Evaluation of *F*_sph_

Although we determine the spherical solutions numerically, we first present two simple theoretical estimates for *F*_sph_*/*(*D* Γ) that serve as useful reference limits. Let *a* denote the radius of the circular contact area between the spherical capsule and the plates. Assuming a constant internal pressure *p*_0_, the force acting on the plates is *F*_sph_ = *πa*^2^*p*_0_. In the cylindrical case, the contact force density is Γ = *p*_0_*W*. Matching the contact geometries by setting 2*a* = *W* (cylindrical contact limit), and using 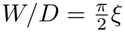 from Eq. (22), we obtain

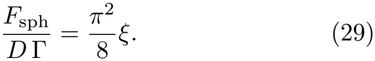

Alternatively, a different estimate of *a* follows from the intersection of an undeformed sphere with a plane at an indentation depth of 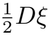. This geometric construction gives 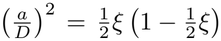 (geometric Hertzian limit [8, 16, 36]), and therefore

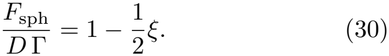

Fig. 4A shows the simulation results for *F*_sph_*/*(*D* Γ). The curves exhibit only a weak dependence on 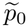, and the theoretical estimates in Eqs. (29) and (30) bracket the simulation results over the range *ξ <* 0.5. In particular, the inequality 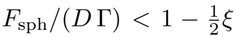 holds in the small-compression limit because stress regularization near the contact perimeter reduces the actual contact area below the geometric Hertzian estimate [8, 36]. The simulation results are well approximated by the bounded exponential function

**FIG. 4:**
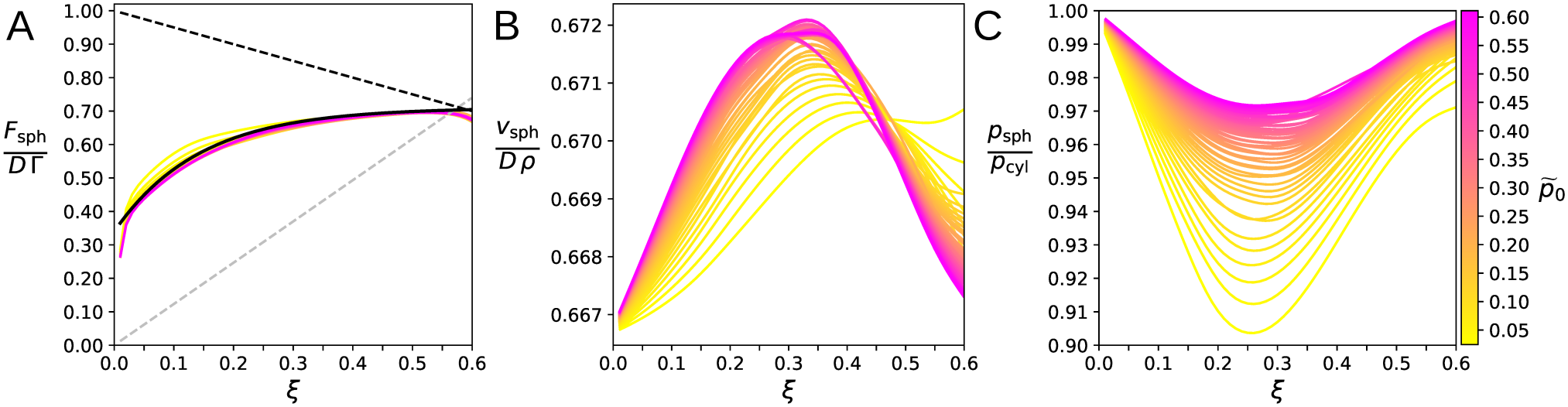
Numerical simulation of parallel-plate compression of pressurized spherical capsules. Simulations were performed with the elastic model of Fig. 2 (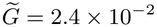 and *ϵ*_*c*_ = 0.04), for 49 different values of 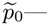 reverse *spring* colormap shown in (C)— in the range 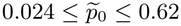 **(A)** Compression force ratio 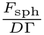 versus *ξ*. The gray dashed line represents the cylindrical contact limit 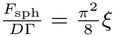, Eq. (29). The black dashed line represents the geometric Hertzian limit 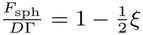, Eq. (30). The solid black line represents the bounded exponential curve *φ*(*ξ*) = *φ* + *k* [1 − exp(−*τξ*)], with *φ* = 0.34, *k* = 0.37 and *τ* = 7.00. **(B)** Volume ratio 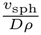 versus *ξ*. **(C)** Pressure ratio 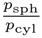 versus *ξ*.

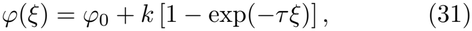

where *φ*_0_ = 0.34, *k* = 0.37, and *τ* = 7.00. The resulting curve is represented by the solid black line in Fig. 4A. This approximation avoids the need for numerical interpolation and expresses the renormalization factor in Eq. (28) as

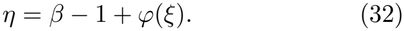

We use this expression directly to compute the master curve *Q* (27), shown in the inset of Fig. 2. To further validate the force decomposition (26), we demonstrate that using *η* from Eq. (32) produces a master curve under different conditions, as shown in Fig. A.4 (Appendix G 1). Although the functional form of *φ*(*ξ*) is unchanged, its parameters (*φ*_0_, *k*, and *τ*) depend slightly on the elastic model. This dependence is characterized in Appendix F 2, where Table I lists the fitted parameters for different values of the characteristic strain for hardening *ϵ*_*c*_.

**TABLE 1:**
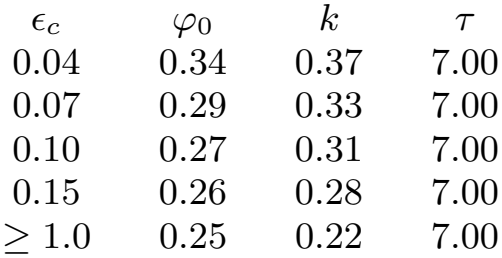
Bounded exponential parameters. Summary of the parameters of the bounded exponential function *φ*(*ξ*) = *φ*_0_ + *k* [1 − exp(−*τξ*)] used to approximate 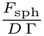 for different values of *ϵ*_*c*_. Since the strain typically remains below 0.30 during compression, values of *ϵ*_*c*_ *≥* 1.0 correspond to a practically linear elastic regime over the range of deformations considered.

| ϵ<sub>c</sub> | φ<sub>0</sub> | *k* | τ |
|---|---|---|---|
| 0.04 | 0.34 | 0.37 | 7.00 |
| 0.07 | 0.29 | 0.33 | 7.00 |
| 0.10 | 0.27 | 0.31 | 7.00 |
| 0.15 | 0.26 | 0.28 | 7.00 |
| ≥ 1.0 | 0.25 | 0.22 | 7.00 |

### C. Contact Area Model

Because the osmotic pressure depends on the total cell volume, the volume decomposition in Eq. (26) can be substituted into Eq. (4) to obtain an approximation for the pressure (Appendix E). As shown in Eq. (E4), this pressure is a weighted sum of the pressures *p*_sph_ and *p*_cyl_ associated with the spherical and cylindrical solutions, respectively, each determined by its corresponding volume contribution, *v*_sph_ or (*β* − 1)*Dρ*.

The construction in Eq. (26) thus defines both the force and the pressure, and therefore specifies the total contact area directly through their ratio, *A*_*c*_ = *F/p*, introduced in Eq. (13), as the renormalized form

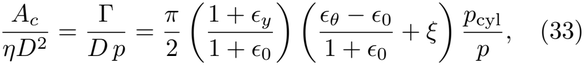

where *p* is given by Eq. (E4), and *ϵ*_*y*_ and *ϵ*_*θ*_ are the strains of the cylindrical solution (section IV B).

Fig. 4B shows that the volume ratio *v*_sph_*/*(*D ρ*) remains nearly constant over a broad range of 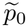, indicating that the relative volume changes of the cylindrical and spherical solutions are approximately the same. Consistent with this observation, Fig. 4C shows that *p*_cyl_ and *p*_sph_ are also similar. Even at the smallest simulated value of 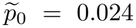, corresponding to a turgor pressure of 0.12 atm for an external osmolality of 200 mOsm/L, the ratio *p*_sph_*/p*_cyl_ remains above 0.92, while at larger turgor pressures it remains close to unity. Under these conditions, using *p* ≈ *p*_cyl_ and *ϵ*_*y*_ ≈ *ϵ*_*θ*_ ≈ *ϵ*_0_, the contact area relation (33) reduces to

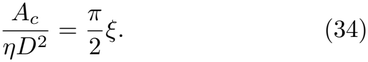

The simulation data in Fig. 4 correspond to the highly nonlinear elastic model considered for *E. coli* (*ϵ*_*c*_ = 0.04). For larger values of *ϵ*_*c*_, however, the behavior of the spherical solution becomes more complex. As shown in Appendix F 2, the ratio *v*_sph_*/*(*D ρ*) increases with *ξ*, with this trend becoming more pronounced at larger values of 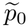 (Fig. A.3A2, B2, C2). This increase is accompanied by a systematic decrease in *p*_sph_*/p*_cyl_ below unity (Fig. A.3A3, B3, C3). In the linear elastic limit, *p*_sph_*/p*_cyl_ reaches ≈0.70 at *ξ* = 0.50 (Fig. A.3A3). Importantly, the force–volume decomposition in Eq. (26) does not require the assumption *p*_sph_ ≈ *p*_cyl_ and remains valid more generally, as demonstrated in Appendix G. Furthermore, as explained in Appendix E, for the average cell aspect ratio (*β* = 4), the full relative pressure *p/p*_cyl_ exhibits substantially weaker variations than *p*_sph_*/p*_cyl_. For *ξ* ≲ 0.30, the simulation data indicate that Eq. (34) remains a good approximation across the elastic models and control parameters considered. In Appendix G 3 and H, we assess the accuracy of the general contact-area model (33) by comparing its predictions with numerical simulations over a range of aspect ratios, elastic models, and control parameters. Across all cases considered, Eq. (33) remains in close agreement with the simulation results (Figs. A.6, A.9, and A.11).

## VI. FINITE-BEAD CURVATURE AND EFFECTIVE GEOMETRY

When compression is applied using a large spherical bead, rather than a parallel plate, the contact on a cylindrical body is not homogeneous (see Fig. 5, comparing the contact areas generated by a large bead and a plate). This spatial variation reflects the mismatch in Gaussian curvature between the two contacting surfaces. In particular, the contact length *W* varies along the longitudinal direction, as illustrated schematically in Fig. 6. We now incorporate this geometric effect into the spherocylindrical model developed in the previous section.

**FIG. 5:**
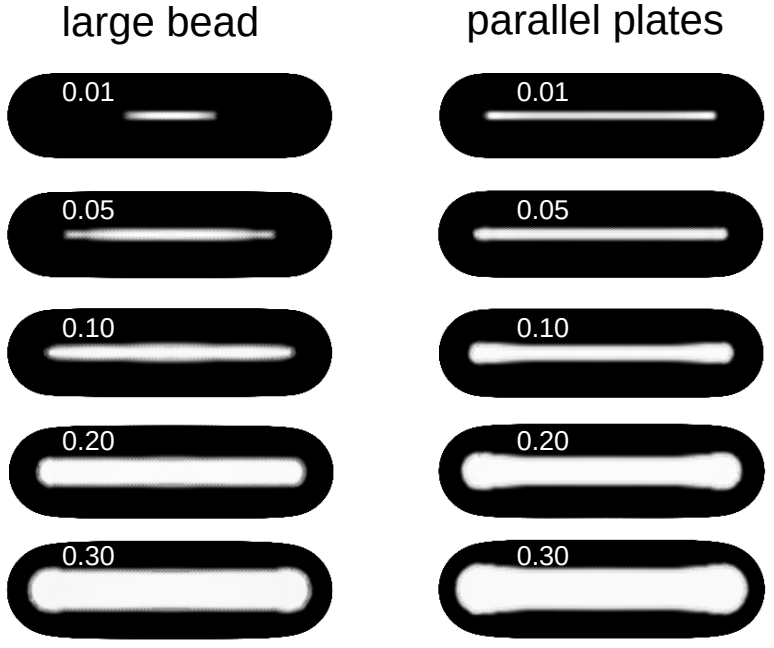
Contact region comparison. Simulation results 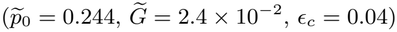 showing the contact regions obtained with large beads (*R/D* = 34) and parallel plates at different relative compression amplitudes (*ξ* = 0.01, 0.05, 0.10, 0.20, 0.30). The images show the magnitude of the contact force exerted on the capsules using a reversed grayscale color map.

**FIG. 6:**
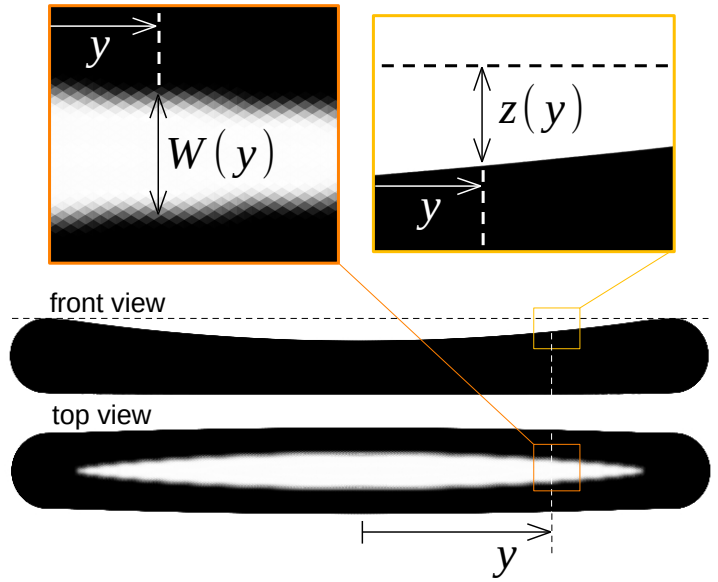
Parametrization of the contact profile. The shapes and configurations shown in the diagrams correspond to simulation outputs 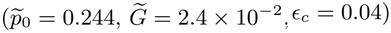 of a capsule with aspect ratio *β* = 10 compressed by a large spherical bead (*R/D* = 34) at *ξ* = 0.30.

The points (*x, y, z*) on the spherical bead defining the contact surface satisfy

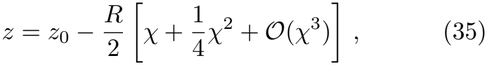

where 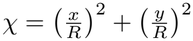 and *z*_*0*_ is the indentation depth at the center of the contact. The *y*-axis is parallel to the long axis of the bacterium, while the *x*-axis lies in the tangent plane of the bead at (*x, y*) = (0, 0) and is perpendicular to both the *y*-axis and the direction of bead displacement.

The contact region remains narrow along the *x*-axis, with |*x*| ≤ *W* and *W* = *W* (*y*) at most of order *D*. By contrast, the longitudinal coordinate *y* can extend over the entire cylindrical body. Consequently, the indentation profile is dominated by its dependence on *y*, justifying the approximation *χ* ≈ (*y/R*)^2^, which yields

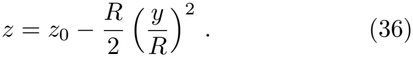

To express the indentation profile as a function of the compression amplitude, we assume *z*_0_ = *D ξ*, consistent with the strong confinement imposed by the bead; that is, there is no geometrical gap between the highest point of the capsule and the height at which the contact profile vanishes. Eq. (36) thus shows that the bead remains in contact over a longitudinal distance *l* given by

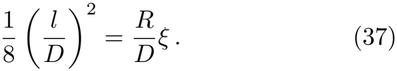

As discussed in the previous section, the response of the capsule can be represented by an equivalent cylinder of aspect ratio *η*, Eq. (28). The longitudinal contact length produced by the bead therefore becomes comparable to the entire equivalent cylinder when *l/D* = *η*. This condition defines a threshold compression amplitude, *ξ*_*t*_, above which the bead contact spans the entire equivalent cylinder. For sufficiently large bead radius, we expect *ξ*_*t*_ ≪ 1. Moreover, using (32), for small compression we have *η*^2^ ≈ (*β* −1 + *φ*_0_) (*β* − 1 + *φ*_0_ + 2*kτξ*), which yields

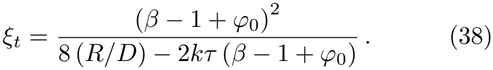

With *φ*_0_ = 0.34, *k* = 0.37, *τ* = 7.00 (solid black line in Fig. 4A) and the bead radius used in the simulations, *R/D* = 34, Eq. (38) predicts *ξ*_*t*_ ≈ 0.05 for *β* = 4.0 and *ξ*_*t*_ ≈ 0.13 for *β* = 6.0. The dashed line in the inset of Fig. 7 shows the approximation given by Eq. (38), while the red line represents the exact numerical evaluation of *ξ*_*t*_ = *ξ*_*t*_(*β*) using the original definition of *η*, Eq. (28), and spline interpolations to represent *F*_sph_*/*(*D* Γ) from the simulation data in Fig. 4A. The close agreement between the two curves confirms that Eq. (38) provides a good approximation over the range of aspect ratios relevant to the present analysis.

**FIG. 7:**
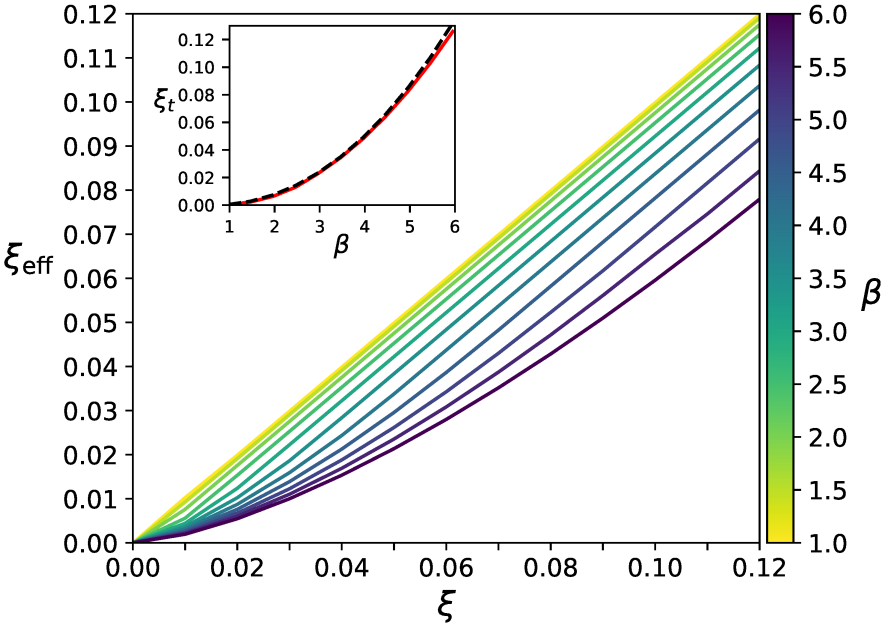
Effective relative compression curves. *ξ*_eff_ versus *ξ* for capsules spanning the aspect-ratio range 1.0≤ *β* ≤ 6.0 (reverse *viridis* colormap) compressed by the same bead (*R/D* = 34). The curves were obtained from Eq. (48) with Δ*ξ*_*t*_ = 0.005 and using *η* from Eq. (32) with *φ*_0_ = 0.34, *k* = 0.37 and *τ* = 7.00. Inset: Corresponding threshold displacement *ξ*_*t*_ as a function of the aspect ratio *β*; the red line is computed using *η* from Eq. (28) and taking 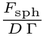 as an interpolation function of the simulation data presented in Fig. 4A, while the dashed line is obtained directly from Eq. (38).

To connect the bead-compression problem to the equivalent parallel-plate description, we define the effective relative compression amplitude as the average indentation over the normalized length *η* of the equivalent cylinder:

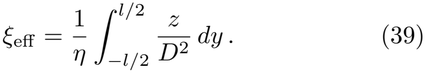

This definition provides a mapping between the finitebead geometry and the parallel-plate compression problem. A convenient way to illustrate this correspondence is through the contact area

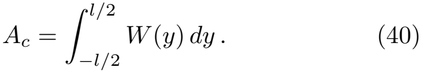

Assuming that each local cross section follows the same flattening kinematics as the cylindrical model, Eq. (22) can be applied locally, giving

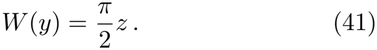

Substituting this relation into Eq. (40) gives

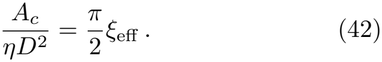

This recovers the parallel-plate result in Eq. (34) with the replacement *ξ → ξ*_eff_.

Using the parabolic bead profile, Eq. (39) yields

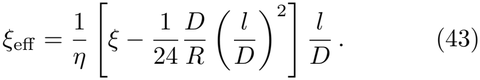

For *ξ > ξ*_*t*_, we set *l/D* = *η*, and Eq. (43) reduces to

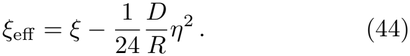

Conversely, for *ξ < ξ*_*t*_, substituting Eq. (37) into Eq. (43) gives

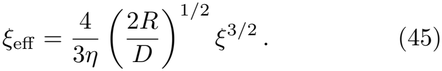

This result implies that the contact-area relation, Eq. (42), becomes independent of *β*, consistent with a contact region localized around the capsule center. In the small-indentation regime, volume variations are negligible and the pressure therefore remains approximately constant. Consequently, the normalized force scales as

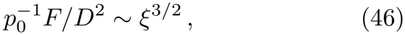

recovering the characteristic Hertzian scaling of localized contact mechanics [16].

Assuming that the transition between the two regimes occurs over a narrow compression interval Δ*ξ*_*t*_ centered at *ξ*_*t*_, we interpolate between them using the sigmoidal switching function

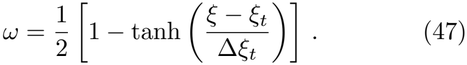

The effective compression is then given by

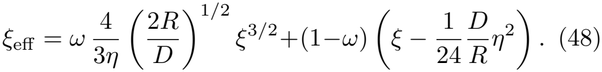

Eq. (48) provides the full expression used to map *ξ* to *ξ*_eff_. The aspect ratio *β* enters this mapping only through *η*. Fig. 7 shows the resulting curves for different values of *β*, using *η* as given in Eq. (32). These curves define the horizontal axes of the insets in Fig. 2B,E.

The importance of this mapping is evident in Fig. A.7 (Appendix H), where the force master curve is obtained only when the data are expressed in terms of *ξ*_eff_. The same mapping also recovers the volume master curve, as shown in Fig. A.8, and yields consistent predictions for the contact area, as demonstrated in Fig. A.9. To further validate the method across different bead radii, we perform simulations spanning 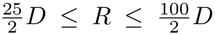. As shown in Appendix H, Figs. A.10 and A.11, the force and contact-area curves follow the same master curves across bead radii after applying the effective-compression mapping and the corresponding force and area renormalizations. Together, these results support the use of the effective compression in the force and volume decomposition given by Eq. (26) and demonstrate the applicability of the finite-bead curvature correction over the range of bead radii investigated.

## VII. DISCUSSION AND CONCLUSIONS

We have developed a theoretical model of the largebead AFM compression of pressurized rod-shaped bacteria. The framework combines a tension-dominated description of the cell envelope with an approximation for spherocylinders of arbitrary aspect ratio based on the spherical and cylindrical limiting solutions, together with a correction for the curvature of the AFM probe.

The model captures the contribution of the hemispherical caps through a renormalized length scale, yielding an equivalent cylindrical representation. Probe curvature produces a spatially heterogeneous contact region that is particularly relevant at small indentation depths. We account for this effect through the effective compression amplitude *ξ*_eff_, which maps the bead-compression geometry onto an equivalent parallel-plate compression problem. Numerical simulations further show that this mapping, combined with the appropriate force, volume, and contact-area renormalizations, provides a consistent description across a broad range of aspect ratios, elastic models, control parameters, and bead radii.

Beyond predicting the force response, the framework provides a contact area model (Section V C) that relates the instantaneous force to the internal pressure through *p* = *F/A*_*c*_. Together with the model prediction for *A*_*c*_, this relation allows the pressure to be inferred throughout the compression process from force measurements. Moreover, because in the definition of the master curve *Q*, given by Eq. (27), the initial turgor pressure *p*_0_ acts as a global scaling factor, it can be estimated from measurements spanning a broad range of compression amplitudes, rather than being restricted to the small-compression regime. This flexibility may improve the robustness of the estimate.

The model further predicts that the geometric observables *H*_*z*_, *H*_*y*_, and *H*_*x*_ remain strongly coupled through simple geometric relations (Fig. 2). Because these quantities are directly accessible through microscopy, experimental measurements of their relationships provide an independent test of the modeling framework [12].

The model is expected to be applicable across a broad range of bacterial envelope architectures. In particular, many Gram-negative and Gram-positive bacteria lie within the tension-dominated regime because the characteristic bending length (*B/p*)^1*/*3^ remains much smaller than the bacterial diameter under physiological conditions. Nevertheless, the model assumes quasistatic elastic equilibrium and neglects time-dependent viscoelastic relaxation, adhesive interactions, and frictional effects between the AFM bead and the cell surface. Future extensions could incorporate viscoelastic dissipation, localized envelope defects, and active cell-wall remodeling during loading.

## Appendix

### Appendix A: Elastic Properties of the Bacterial Capsule

#### 1. Elastic Energy and Response Function

Following the nonlinear description of the *E. coli* cell wall developed in Ref. [1], the spherocylindrical capsules with aspect ratios *β* are modeled as triangular networks composed of nonlinear springs with average dimensionless resting length *ã* and average dimensionless resting spring constant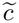. The total number of springs *N* ~ *β/ã*^2^ is typically larger than 10^4^ (see Fig. A.1 for visualization of a capsule of aspect ratio *β* = 2). The elastic energy of the capsule is

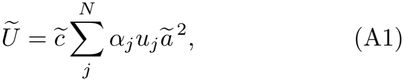

where *u*_*j*_ = *u*_*j*_(*ϵ*) is the reduced elastic energy of spring *j* under intrinsic strain *ϵ*, while the coefficients *α*_*j*_ are weight factors that allow us to specify elastic anisotropy.

All springs share the same response function

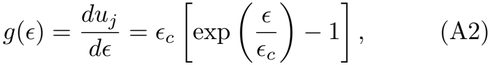

where *ϵ*_*c*_ is the characteristic strain for hardening. The typical value found for *E. coli* in Ref. [1] is *ϵ*_*c*_ = *≫* 0.04. Linear springs are recovered in the limit *ϵ*_*c*_ 1. The initial value *g*_0_ = *g*(*ϵ*_0_) was defined in Eq. (10).

Because the *E. coli* cell wall is extremely thin and the experimentally relevant configurations are tension dominated, the elastic energy does not include an explicit bending contribution. In Sec. IV A, we presented an analytical treatment for long capsules that incorporates bending stiffness and showed that the bending contribution is negligible for both Gram-negative *E. coli* and Gram-positive *B. subtilis* (Sec. IV B).

#### 2. Architecture and Elastic Anisotropy

The elastic anisotropy of the capsule is established through the energy weights *α*_*j*_ introduced in Eq. (A1). They are assigned such that the pre-strain is isotropic [1], independent of the turgor pressure.

As indicated in Fig. A.1, the local arrangement of springs at the hemispherical end caps and the cylindrical region are different. At the caps, the network is constructed with a large number of randomly distributed irregularities and a constant energy weight

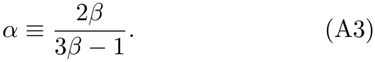

**FIG. A.1:**
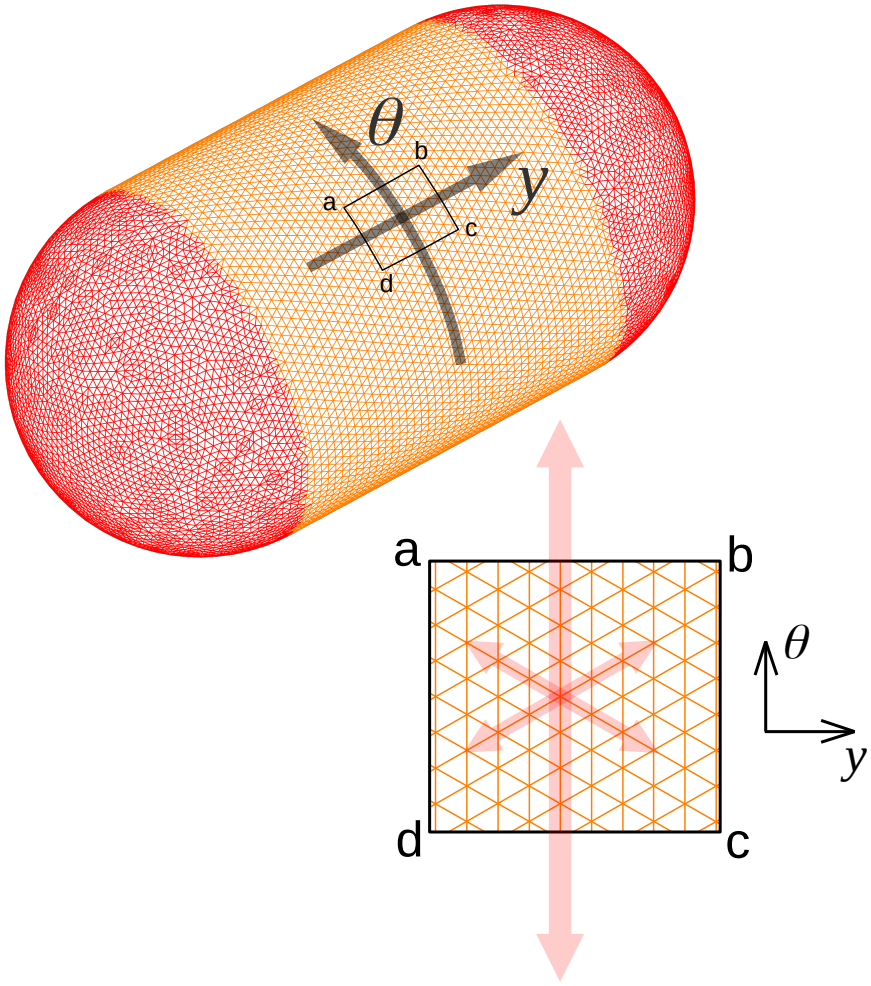
Capsule architecture. Depiction of a capsule used in the simulations (aspect ratio *β* = 2.0). The capsule surface is represented by a triangular spring network, with each edge corresponding to a spring characterized by the response function *g*(*ϵ*), defined in Eq. (A2). The orange and red colors distinguish the cylindrical body from the hemispherical caps. In the cylindrical region, the network has a hexagonal arrangement. The patch abcd shows a close-up of this region: the vertical edges are aligned with the hoop direction (*θ*) and have an elastic energy weight 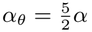, whereas the diagonal edges have weight *α*_*d*_ = *α*. Consequently, the hoop-aligned springs are 2.5 times stiffer than the diagonal springs. This anisotropy, illustrated by the relative sizes of the reddish arrows in the patch, ensures an isotropic pre-strain of the capsule, independent of the turgor pressure. In the hemispherical caps, the network includes numerous randomly distributed irregularities, and all springs have the same energy weight *α*.

It can be inferred from Eq. (B5) that the average elastic constant of the hemispherical caps is 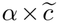, where 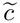 is the global average introduced in Eq. (A1).

In the cylindrical region, the network consists of the hexagonal arrangement of springs described in Ref. [1]. The springs aligned with the hoop direction (*θ*) have energy weights 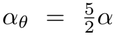, while the diagonal springs have energy weights *α*_*d*_ = *α*. This construction fulfills the isotropic pre-strain condition, with a circumferential Young’s modulus approximately twice the longitudinal one and a uniform shear modulus 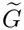 throughout the capsule.

One can show that 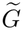 and 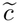 are related through

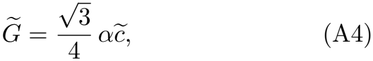

and the Young’s moduli in the hoop and longitudinal direction (*y*) satisfy

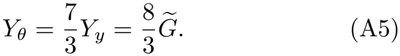

#### 3. Spatial Tension Densities

Eq. (A1) implies that a spring with stiffness 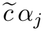 and strain *ϵ* exerts a force proportional to 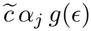. Following Ref. [1], this force can be converted into a spatial tension density,

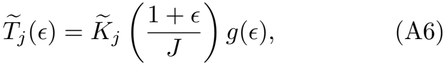

where

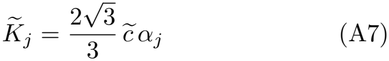

is the dimensionless nominal rigidity at rest, and *J* is the determinant of the deformation gradient tensor. The numerical factor 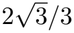 corresponds to the average ratio of the squared spring length to the combined area of the two adjacent triangles sharing that spring. We next use the definition (A6) to derive the stress–strain relationships for the cylindrical part of the capsule, as detailed in Appendix A 4.

#### 4. Stress-Strain Relationships

For the elastic energy defined in (A1), we derive the corresponding constitutive stress–strain relationships, accounting for the elastic anisotropy described in Appendix A 2. These relations are used in the analytical treatment of cylinders in Sec. IV B to compute the curves shown in Fig. 3.

For a deformation defined by the principal strains *ϵ*_*θ*_ and *ϵ*_*y*_—corresponding to the hoop strain and longitudinal strain respectively—the determinant of the associated deformation gradient tensor is

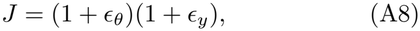

and the spatial tension densities (A6) for the cylindrical region are

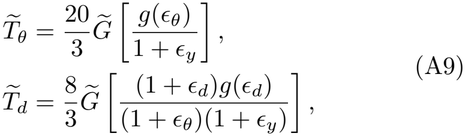

where 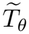 and 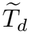 denote the tensions associated with the hoop and diagonal springs, and the diagonal strain *ϵ*_*d*_ satisfies

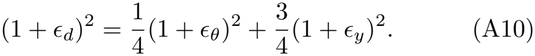

In the continuum limit of the network, the stress–strain relationships for the cylindrical region become [1]

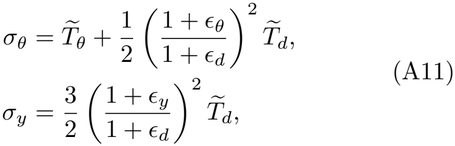

where *σ*_*θ*_ and *σ*_*y*_ are the principal components of the Cauchy stress tensor.

Substituting (A9) into (A11) yields

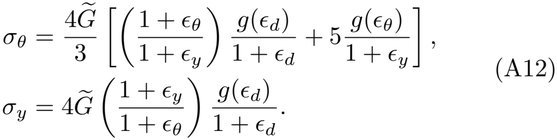

### Appendix B: Shear Modulus Compatibility

The average spring constant 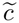, first introduced in Eq. (A1), can be expressed as a function of the initial dimensionless turgor 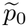 and pre-strain *ϵ*_0_ through the initial static equilibrium condition. By definition, the energy weights satisfy

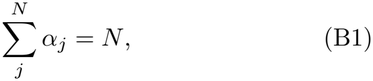

and, because the initial configuration has an isotropic pre-strain, its elastic energy is

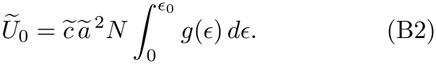

Using the geometrical identity

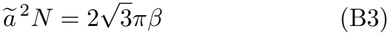

for hexagonal networks, the equilibrium condition

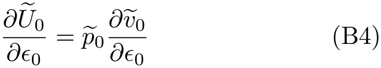

yields

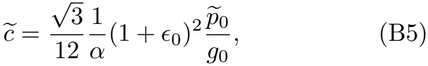

where *g*_0_ = *g*(*ϵ*_0_) is the initial value of the response function (A2). The same expression follows directly from the stress–strain relationships in Eq. (A12) by imposing mechanical equilibrium.

Because the shear modulus is uniform throughout the capsule (see Appendix A 2), Eq. (A4) can be combined with Eq. (B5) to eliminate the dependence on *β*, yielding

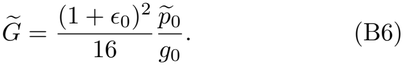

This relation is equivalent to Eq. (9) in the main text.

### Appendix C: Equilibrium Equation for Cylinders

The shape of the cross-section of the cylinder is governed by the *elastica* equations [18]:

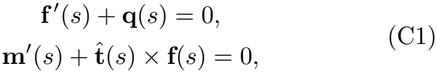

where **q**(*s*) is the external force density, **f**(*s*) is the internal force density—whose tangential component equals the hoop stress *σ*_*θ*_ of the cylinder—, **m**(*s*) is the internal torque, the apostrophes denote derivatives with respect to the arc length coordinate, *s*, and 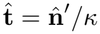 is the tangential direction, where 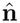 denotes the normal vector and *κ* = *κ*(*s*) is the local curvature.

The external force density is assumed normal to the surface of the cylinder,

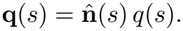

With 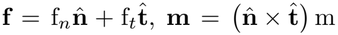, and 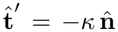, equations (C1) lead to the scalar equations

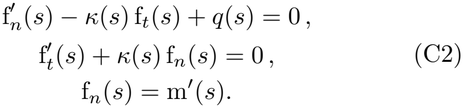

Using the last expression to eliminate f_*n*_, we obtain

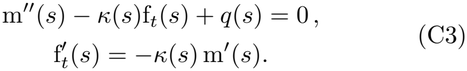

Although bending contributions were neglected in the definition of the elastic energy of the capsule (A1), it is useful for the present derivation to retain a bending stiffness, denoted by *B*. The role of bending effects is examined in the main text, Sec. IV B. The internal torque is then related to the local curvature through

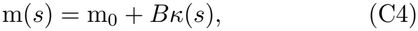

where m_0_ is a constant characterizing the natural curvature of the capsule. Substituting (C4) into the second equation of (C3) and integrating once yields

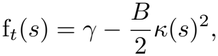

where *γ* is an integration constant equal to the hoop stress within the flat contact regions. Since the total force *pH*_*z*_ exerted by the lateral free portions of the cross-section must be balanced by the tension transmitted through the two contact regions, we obtain

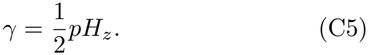

Substituting this result into the first equation of (C3) finally leads to the differential equation governing the cross-sectional shape,

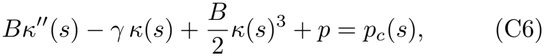

where *p*_*c*_(*s*) is the contact pressure, so in general *q*(*s*) = *p* − *p_c_*(*s*)

### Appendix D: Small-Displacement Analysis

To better understand how the small-compression response depends on 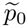 and the constitutive properties of the envelope, we expand the equilibrium equations for the cylindrical part of the capsule, Eqs. (A11) and (19), perturbatively in powers of the relative compression amplitude *ξ*.

The strain components are expanded as

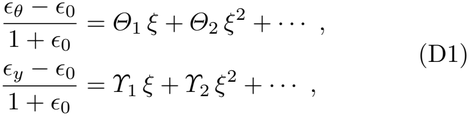

where the coefficients *Θ*_1_, *γ*_1_, *Θ*_2_, *γ*_2_, and higherorder terms depend on 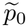, *ϵ*_0_ and the response function *g*(*ϵ*) (A2).

Combining these expansions with the geometrical relations (16), (17) and (21), derived in Section IV B and IV C of the main text, yields

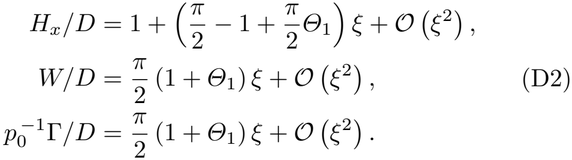

The corresponding relative variations of the envelope area and enclosed volume are

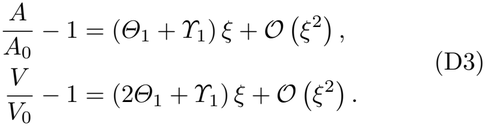

To linear order, the dependence on *ϵ*_0_ and *g*(*ϵ*) enters through the parameter

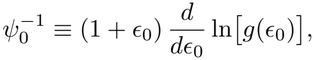

where *ψ*_0_ is the generalized pre-strain [1].

To linear order, combining Eqs. (D1) with Eqs. (A12) and (19) yields

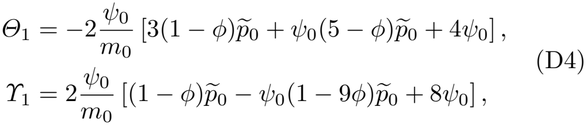

where

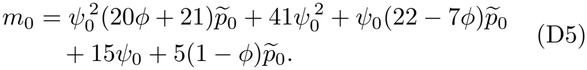

The coefficient *Θ*_1_ is always negative, whereas *γ*_1_ is always positive. Thus, compression reduces the crosssectional perimeter, as reflected by the decrease in *ϵ*_*θ*_ in Eq. (D1), while inducing longitudinal elongation of the cylinder, as reflected by the increase in *ϵ*_*y*_. Both effects are small because *Θ*_1_ and *γ*_1_ are proportional to *ψ*_0_, which is generally small (*ψ*_0_ ≈ 0.03 for *E. coli* [1]).

In the limit 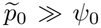, corresponding to strong strain hardening, the pre-strain remains small even at very high turgor, and the first-order coefficients reduce to

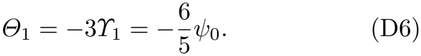

Conversely, in the weak strain-hardening regime, 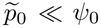, we obtain

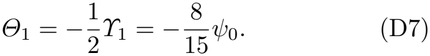

These limiting cases show that the strain variations induced by compression remain parametrically small across a broad range of physiological conditions, explaining why the cross-sectional perimeter remains nearly constant.

### Appendix E: Pressure Decomposition

The pressure is strictly dependent on the volume through the van ‘t Hoff relation given by Eq. (4). Our decomposition of the volume (26) has the form *v* = *v*_cyl_ + *v*_sph_, where *v*_cyl_ = (*β* − 1) *D ρ* is the amount of volume contributed by the cylindrical solution. Similarly, the total volume for which the turgor vanishes, *µ*, can be decomposed as *µ* = *µ*_cyl_ + *µ*_sph_, where *µ*_cyl_ and *µ*_sph_ refer to the contributions of the cylindrical solution and spherical solution such that both solutions retain the same initial turgor. In other words,

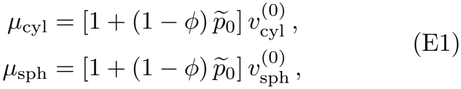

where 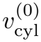 and 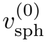 are the associated volumes of the cylindrical and spherical solutions at *ξ* = 0, and 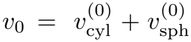. With these definitions, Eq. (4) can be expressed in the form

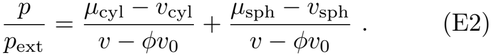

Then, using the volume fractions

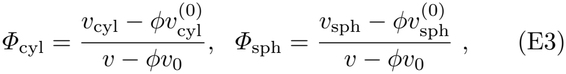

we find the pressure decomposition formula

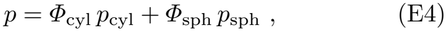

where *p*_cyl_ and *p*_sph_ are the pressures of the cylindrical and spherical solutions,

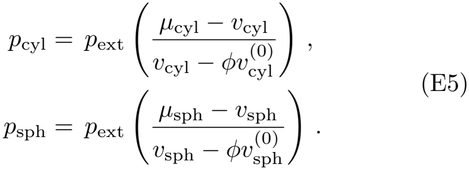

Note that the volume fractions satisfy *Φ*_cyl_ + *Φ*_sph_ = 1, so they act as natural weighting factors for the two asymptotic solutions. Using Eq. (E4), the pressure can be rewritten in the form

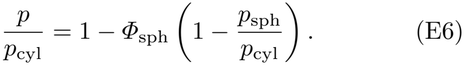

For long capsules, *Φ*_sph_ is typically small, so we expect the ratio 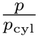 to vary less strongly than 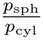. Indeed, for *ϕ* ≪ 1 we find

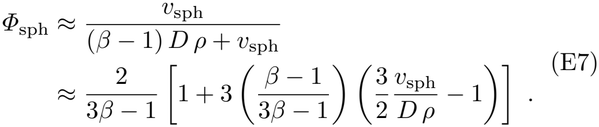

In Fig. A.3, the ratio 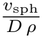 reaches a maximum value of approximately 0.71 at *ξ* = 0.5 for the linear elastic model (Fig. A.3A2). For an average aspect ratio of *β* = 4, Eq. (E7) then gives *Φ*_sph_ ≈ 0.19. Thus, using Eq. (E6) and the lowest observed value of 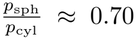 at *ξ* = 0.5 (Fig. A.3A3), we obtain *p* ≈ 0.94 *× p*_cyl_. In this worst-case estimate, the pressure of the spherocylinder therefore differs by less than six percent from the value given by the cylindrical solution.

### Appendix F: Elastic Model Effect

#### 1. Effects on the loading force

The weak dependence of the loading force on the elastic model assumed for the capsule material is confirmed by numerical simulations of the spring networks by *Surface Evolver* [5]. The elastic response can be varied through the characteristic strain for hardening *ϵ*_*c*_, defined in Eq. (A2), with the response approaching the linear elastic limit as *ϵ*_*c*_ increases. In the main text, we use *ϵ*_*c*_ = 0.04, corresponding to the estimate for *E. coli* reported in Ref. [1]. Figure A.2 shows the normalized force as a function of compression amplitude *ξ* for a representative capsule (*β* = 4.0) and different values of *ϵ*_*c*_, including the linear elastic limit *ϵ*_*c*_ *≫* 1. The effect of *ϵ*_*c*_ is negligible for *ξ <* 0.20. At larger compression amplitudes, the force response becomes slightly more nonlinear for smaller *ϵ*_*c*_ as expected. A similar trend is observed for the cylindrical solution developed in Sec. IV B.

#### 2.2 Effects on the hemispherical end-cap corrections

As discussed in Sec. V in the main text, the hemispherical contributions to the force and volume are characterized by the ratios 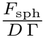 and 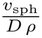. Their dependence on 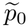 and the elastic model, determined by *ϵ*_*c*_, is shown in Fig. A.3.

**FIG. A.2:**
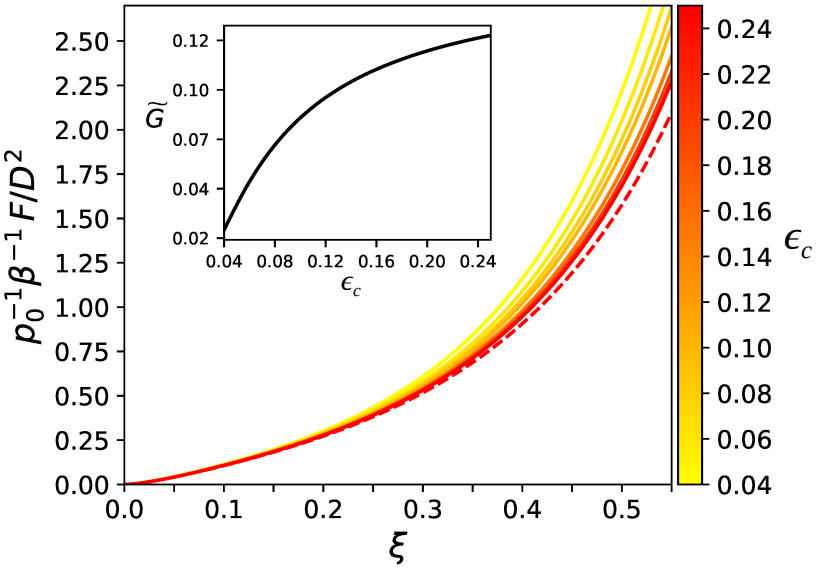
Normalized force for different strain-hardening parameters. Normalized compression force 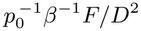 as a function of *ξ* for capsules with *β* = 4 and characteristic hardening strain *ϵ*_*c*_ in the range 0.04 ≤ *ϵ*_*c*_ ≤ 0.25, with colors following the reverse *autumn* colormap. The dashed line represents the linear-spring limit (*ϵ*_*c*_ *≫* 1.0). The curves were obtained from simulations using *Surface Evolver* [5] with 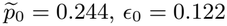, and bead radius *R* = 34 *× D*. The color map also indicates the corresponding dimensionless shear moduli 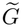, determined by *ϵ*_*c*_ through Eq. (9). The inset shows 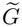 as a function of *ϵ*_*c*_.

Over the broad range of parameters explored, the ratio 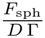 exhibits only a weak dependence on 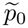 (Fig. A.3A1, B1, C1, D1). We therefore approximate this ratio, following the same strategy as in the main manuscript, using the bounded exponential function *φ*(*ξ*) = *φ*_0_ + *k* [1 − exp(− *τξ*)], introduced in Eq. (31). The force renormalization factor *η*, defined in Eq. (28), is then given by *η* = *β* − 1 + *φ*(*ξ*). Table I summarizes the fitted parameters (*φ*_0_, *k, τ*) for each value of *ϵ*_*c*_ considered.

The dependence of the ratio 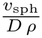 on *ϵ*_*c*_ is more complex. For a strongly nonlinear elastic response, as in the case of *E. coli* considered in the main text (*ϵ*_*c*_ = 0.04), we obtain 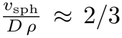 (Fig. A.3D2), which also implies that the pressure of the cylindrical solution, *p*_cyl_, is approximately equivalent to the pressure of the spherical solution, *p*_sph_ (see Fig. A.3D3). These approximations appear to be independent of 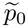, supporting the expression for the contact area *A*_*c*_ given in Eq. (34), where *p* = *p*_cyl_ is used.

For larger values of *ϵ*_*c*_, the ratio 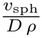 increases with compression amplitude *ξ*, with this trend becoming more pronounced at larger values of 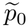 (Fig. A.3A2, B2, C2). This increase is accompanied by a systematic decrease in *p*_sph_*/p*_cyl_ below 1.0 (Fig. A.3A3, B3, C3). In the linear elastic limit, this ratio reaches approximately 0.70 at *ξ* = 0.50 (Fig. A.3A3).

These findings do not affect the applicability of the decomposition of the spherocylindrical response developed in Sec. V. Indeed, Figs. A.4 and A.5 show that Eq. (26) remains valid even in the linear elastic limit. However, when the approximation *p* = *p*_cyl_ does not hold, the pressure must be evaluated using Eq. (E4), and the contact area must then be calculated using Eq. (33).

### Appendix G: Parallel-plate Compression Geometry

We tested the force decomposition *F* = (*β* − 1)Γ *D* + *F*_sph_ and volume decomposition *v* = (*β* − 1)*ρ D* + *v*_sph_, presented in Eq. (26), against numerical simulations using *Surface Evolver* [5] and a parallel-plate compression geometry. We also provide numerical evidence supporting the associated contact area model introduced in Section V C.

#### 1. Force renormalization

Fig. A.4 shows the force normalized by both *p*_0_*βD*^2^ and *p*_0_*ηD*^2^, with 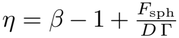 as defined in Eq. (28). We use the approximation 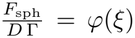, where *φ*(*ξ*) = *φ*_0_ + *k* [1 − exp(− *τξ*)] is the bounded exponential function introduced in Eq. (31), with parameters *φ*_0_, *k*, and *τ* listed in Table I. The renormalized force curves (Fig. A.4A2, B2, C2, D2) collapse onto a common master curve across different aspect ratios *β*, indicating that the force decomposition in Eq. (26) accurately captures the response across a broad range of conditions, including the linear elastic limit (Fig. A.4D2).

#### 2. Volume renormalization

The volume decomposition *v* = (*β* − 1)*ρ D* + *v*_sph_ implies that the relative volume *v/v*_0_, where *v*_0_ is the initial capsule volume used in Eq. (4), can be expressed as

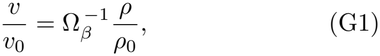

where 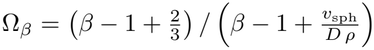 and 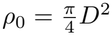 is the value of *ρ* at *ξ* = 0. At zero compression, *ξ* = 0, the volume of the sphere is 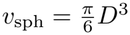, so that 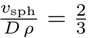 and Ω_*β*_ = 1. Eq. (G1) implies that, if the volume decomposition holds, the product Ω_*β*_ *v/v*_0_ should be independent of the aspect ratio *β*. Thus, Ω_*β*_ acts as a renormalization factor that causes the relative-volume curves to collapse onto a master curve.

For the highly nonlinear elastic response characteristic of *E. coli*, which we focus on in the main manuscript (*ϵ*_*c*_ = 0.04), we find that 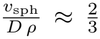 over the full range of compression amplitudes (Fig. A.3D2). Consequently, Ω_*β*_ ≈ 1, and little renormalization is needed to collapse the relative-volume curves. This is confirmed in Fig. A.5, which shows the relative volume obtained from simulations of compression between parallel plates. By contrast, comparison of Fig. A.5D1 and A.5D2 shows that Ω_*β*_ plays an important role in collapsing the curves in the linear elastic limit. These results support the applicability of the volume decomposition in Eq. (26).

#### 3. Contact area renormalization

The simulation data in Fig. A.6 show that, for parallelplate compression, the contact-area curves *A*_*c*_(*ξ*) for capsules with different aspect ratios collapse onto a single curve when normalized by *η D*^2^. This collapse persists over a broad range of control parameters and remains effective even in the linear elastic limit (Fig. A.6D2), where *p*_sph_ is significantly smaller than *p*_cyl_ (see Fig. A.3). The second-row panels (A2, B2, C2, and D2) compare the simulation data with the predictions of the general contact-area model, Eq. (33), evaluated at *β* = 4 (dashed line), and with the linear approximation (dash-dotted lines) given by Eq. (34). The general model closely tracks the simulation data across the cases considered, whereas the linear approximation provides a useful description over the range of relatively small compression amplitudes, *ξ* ≲ 0.30. These comparisons support the use of Eq. (33) as a general approximation for the contact area, including when the spherical and cylindrical pressures differ appreciably.

### Appendix H: Large-Bead Compression Geometry

For compression with a large bead, collapsing the force–compression curves across different aspect ratios requires transforming the compression amplitude according to *ξ → ξ*_eff_(*ξ*), as described in Sec. VI and defined by Eq. (48). Figures A.7, A.8, and A.9 show the resulting collapse of the renormalized force, 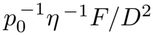, the renormalized volume, Ω_*β*_ *v/v*_0_ (defined in Appendix G 2), and the renormalized contact area, *η* ^−1^*A*_*c*_*/D*^2^, respectively. These results are obtained from *Surface Evolver* simulations [5] spanning a range of external pressures and degrees of elastic nonlinearity, including the linear elastic limit.

To further test the finite-bead curvature correction, we perform simulations for four different bead radii: *R/D* = 25*/*2, 50*/*2, 75*/*2, and 100*/*2. As shown in Fig. A.10, the force–compression curves collapse onto the same master curve after applying the effective-compression transformation and force renormalization. This agreement indicates that the model accounts for the influence of bead radius on the force–compression relationship over the range investigated. The corresponding contact-area curves are shown in Fig. A.11. When normalized by *ηD*^2^ and plotted against *ξ*_eff_, they also collapse closely onto a common curve, showing that the effective-compression transformation accounts for the influence of bead radius on the contact area, as expected.

**FIG. A.3:**
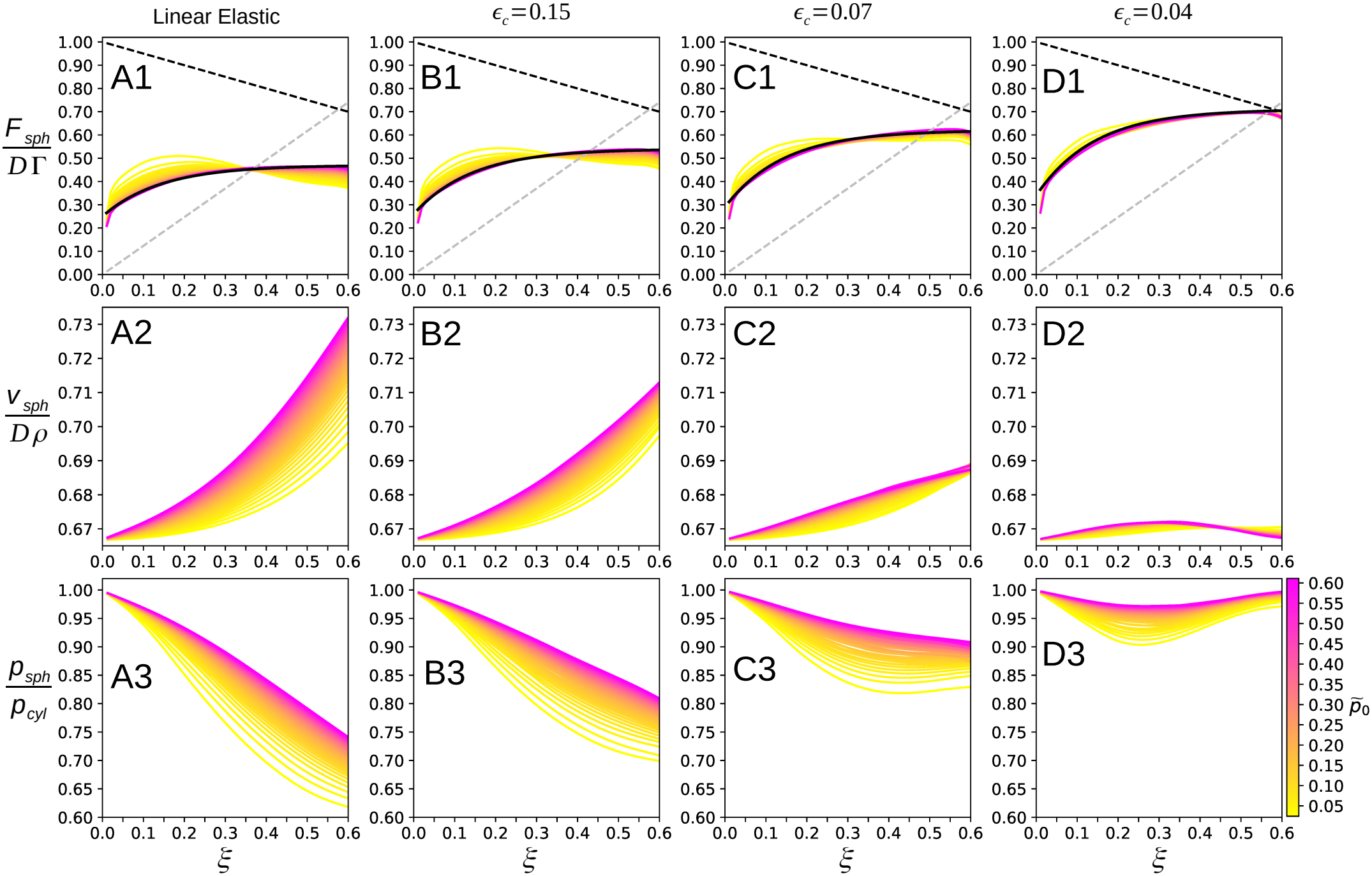
Numerical simulation of parallel-plate compression of pressurized spherical capsules. Simulations were performed using *Surface Evolver* [5] with the same 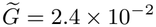 used in Fig. 4, but changing the value of *ϵ*_*c*_ to visualize the effect of the non-linearity of the elastic model. The line colors indicate 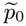, increasing from yellow to pink according to the reverse *spring* colormap used in Fig. 4, over the range 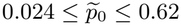 (color bar in the lower-right corner). Columns A–D correspond to a linear elastic model (*ϵ*_*c*_ *≫* 1) and nonlinear elastic models with *ϵ*_*c*_ = 0.15, *ϵ*_*c*_ = 0.07, and *ϵ*_*c*_ = 0.04, respectively, as indicated above the first row. **(A1, B1, C1, D1)** Compression force ratio 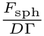 versus *ξ*. The gray dashed line represents the cylindrical contact limit 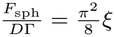, Eq. (29), while the black dashed line represents the geometrical Hertzian limit 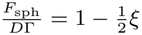, Eq. (30). The solid black line represents the bounded exponential curve (31), *φ* = *φ*_0_ + *k* [1 − exp(−*τξ*)], with *τ* = 7.00 and (*φ*_0_, *k*) = (0.25, 0.22) in A1, (*φ*_0_, *k*) = (0.26, 0.28) in B1, (*φ*_0_, *k*) = (0.29, 0.33) in C1, and (*φ*_0_, *k*) = (0.34, 0.37) in D1. **(A2, B2, C2, D2)** Volume ratio 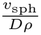 versus *ξ*. **(A3, B3, C3, D3)** Pressure ratio 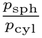 versus *ξ*.

**FIG. A.4:**
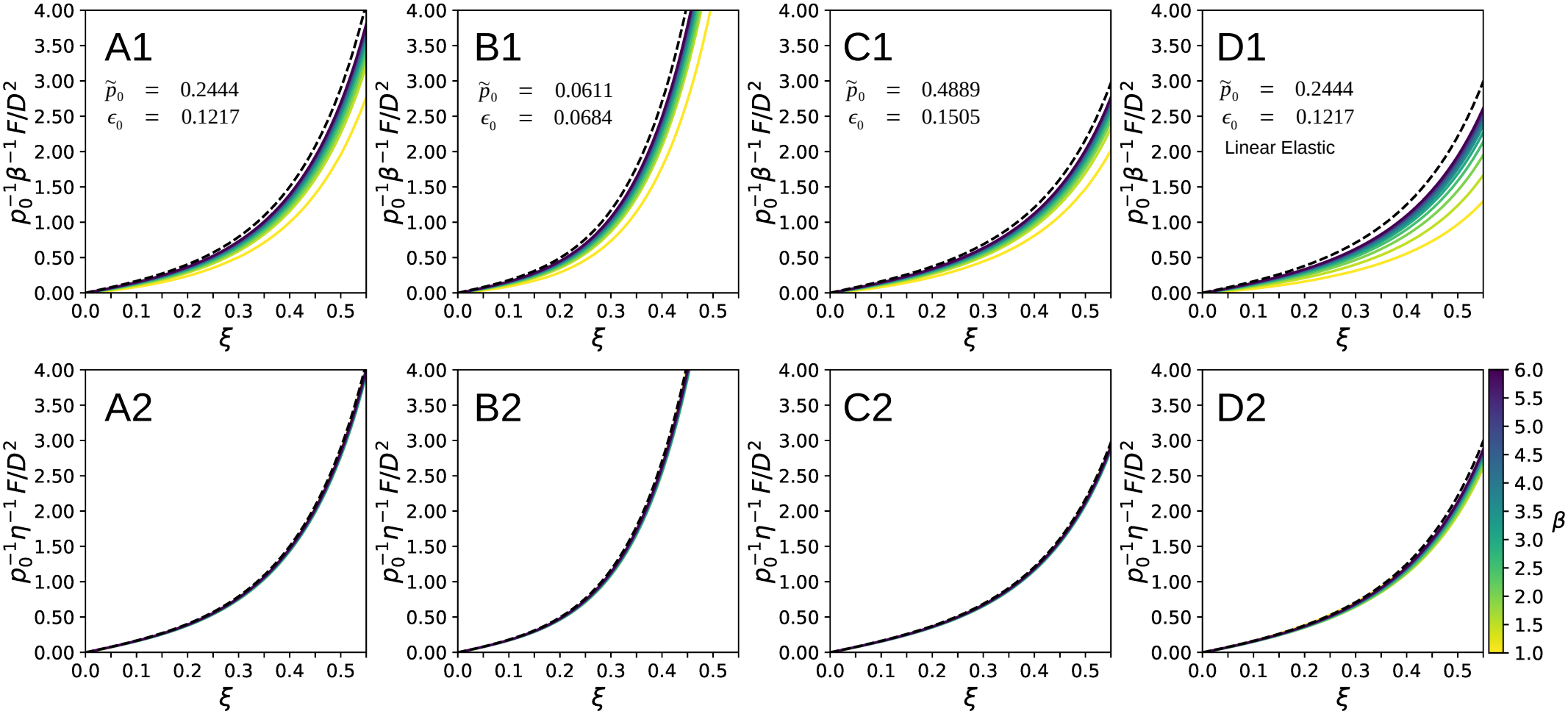
Renormalization of the loading force for parallel-plate compression. Data obtained from simulations using *Surface Evolver* [5] with parallel plates and aspect ratios in the range 1 ≤ *β* ≤ 6. The color of the solid lines indicates *β*, increasing from yellow to blue according to the color map (reverse *viridis*) used in Figs. 2B and 7 (color bar in the lower-right corner). The dashed line corresponds to the normalized force density 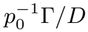 obtained for cylinders in the parallel-plate limit, Eq. (17); in this limit, the curves are unaffected by the renormalizations. Each column (A, B, C, D) corresponds to a different set of control parameters. The values of 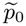 are indicated in the first-row plots: 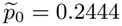 for (A, D), 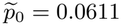 for (B), and 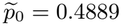 for (C). The first three columns **(A, B, C)** use the same nonlinear elastic model, with 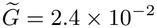 and *ϵ*_*c*_ = 0.04, whereas the last column **(D)** uses a linear elastic model, with 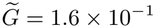 and *ϵ*_*c*_ *→ ∞*. The corresponding pre-strain *ϵ*_0_ is indicated below 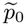 in each first-row plot. The first row **(A1, B1, C1, D1)** shows the force normalized by *p*_0_ *β D*^2^ as a function of the compression amplitude *ξ*. The second row **(A2, B2, C2, D2)** shows the force renormalized by *p*_0_ *η D*^2^, where *η* is given by Eq. (32), with *φ*_0_ = 0.34, *k* = 0.37, and *τ* = 7.00 for (A, B, C), and *φ*_0_ = 0.25, *k* = 0.22, and *τ* = 7.00 for (D), consistent with the parameters provided in Table I.

**FIG. A.5:**
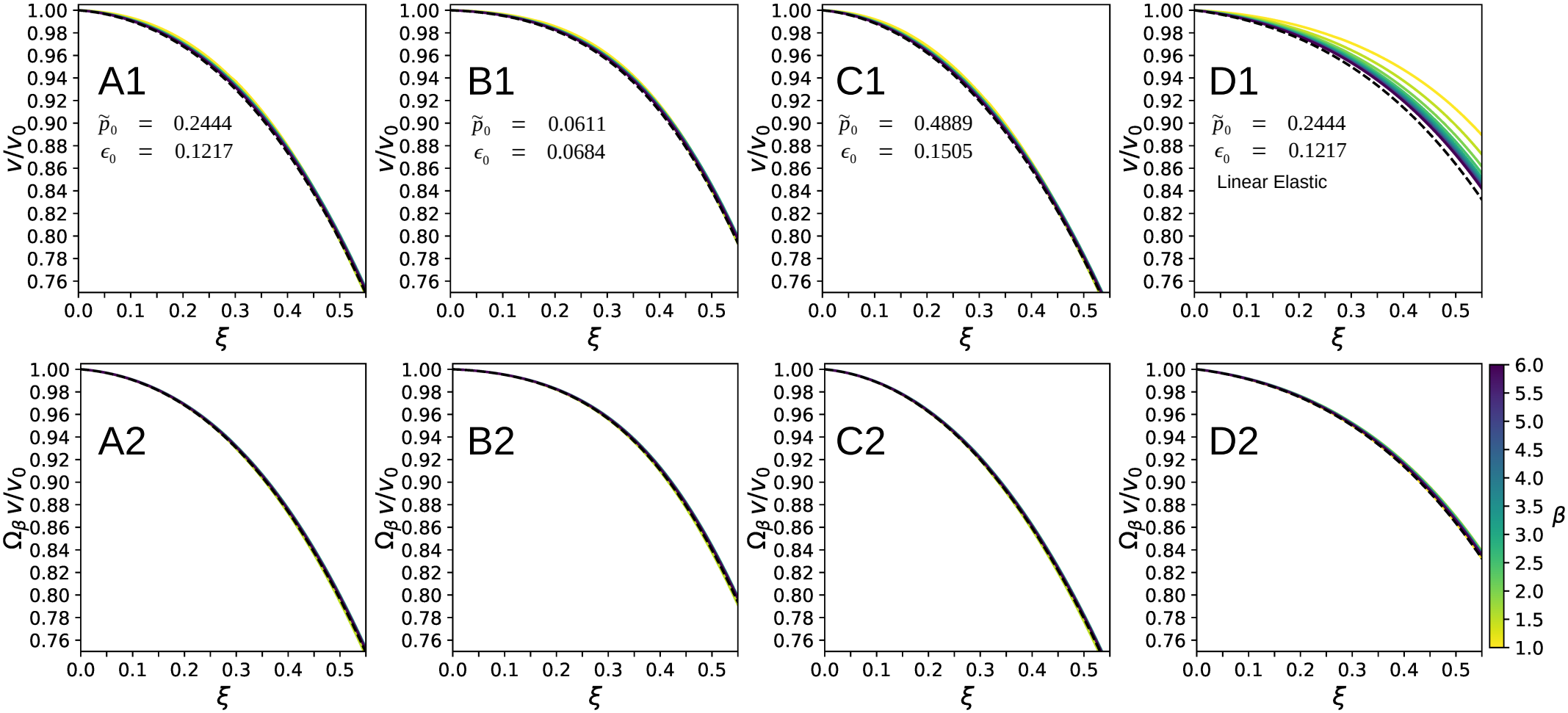
Renormalization of the volume for parallel-plate compression. Data obtained from simulations using *Surface Evolver* [5] with parallel plates and aspect ratios in the range 1 ≤ *β* ≤ 6. The color of the solid lines indicates *β*, increasing from yellow to blue according to the color map (reverse *viridis*) used in Figs. 2B and 7 (color bar in the lower-right corner). The dashed line corresponds to the relative volume density *ρ/ρ*_0_, with 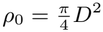, obtained for cylinders in the parallel-plate limit, Eq. (18); in this limit, the curves are unaffected by the renormalizations. Each column (A, B, C, D) corresponds to a different set of control parameters. The values of 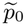 are indicated in the first-row plots: 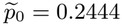 for (A, D), 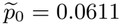 for (B), and 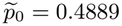 for (C). The first three columns **(A, B, C)** use the same nonlinear elastic model, with 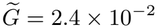 and *ϵ*_*c*_ = 0.04, whereas the last column **(D)** uses a linear elastic model, with 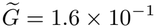 and *ϵ*_*c*_*→ ∞*. The corresponding pre-strain *ϵ*_0_ is indicated below 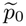 in each first-row plot. The first row **(A1, B1, C1, D1)** shows the relative volume *v/v*_0_ as a function of the compression amplitude *ξ*. The second row **(A2, B2, C2, D2)** shows the relative volume multiplied by the renormalization factor 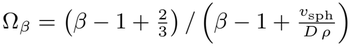.

**FIG. A.6:**
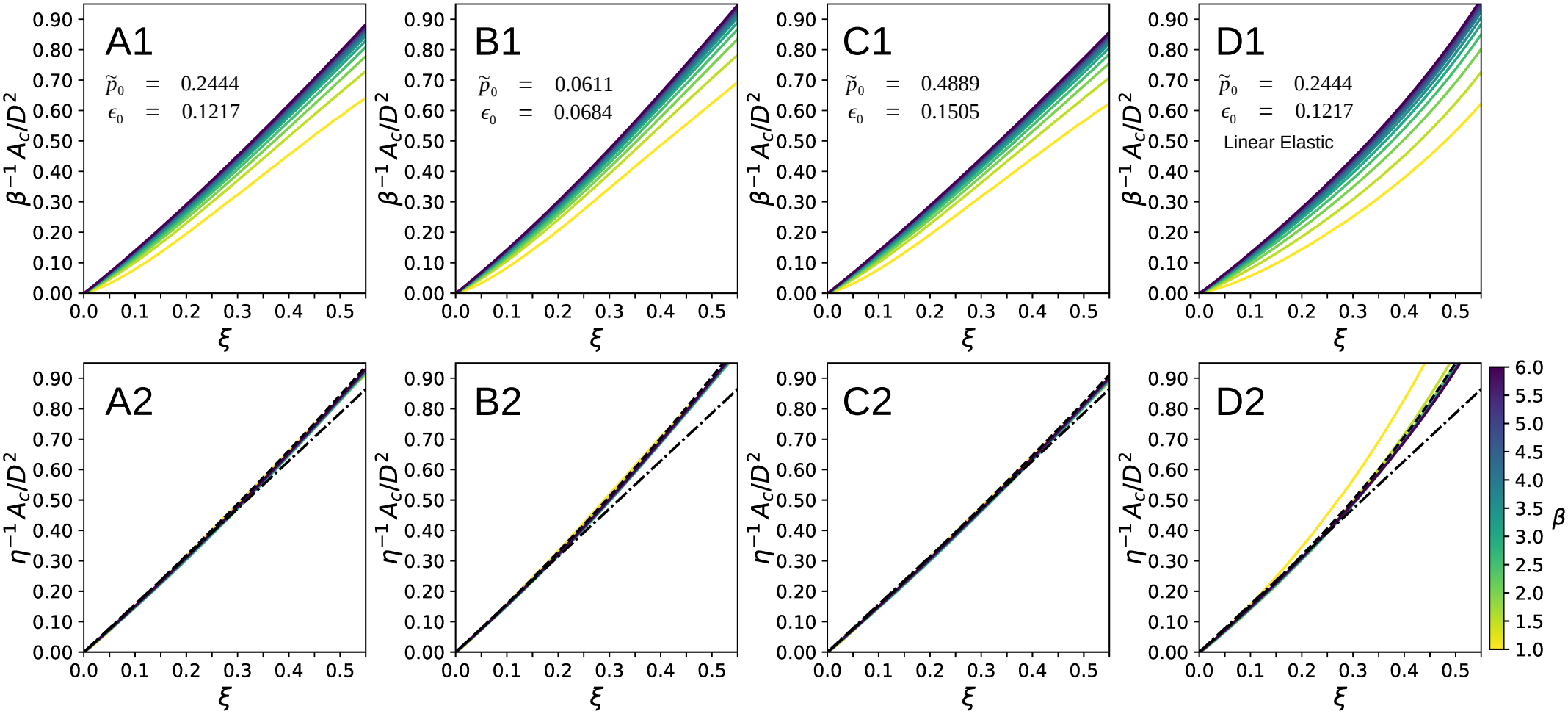
Contact area for parallel-plate compression. Data obtained from simulations using *Surface Evolver* [5] with parallel plates and aspect ratios in the range 1 ≤ *β*≤ 6. The color of the solid lines indicates *β*, increasing from yellow to blue according to the color map (reverse *viridis*) used in Figs. 2B and 7 (color bar in the lower-right corner). Each column (A, B, C, D) corresponds to a different set of control parameters. The values of 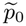 are indicated in the first-row plots: 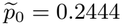 for (A, D), 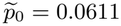 for (B), and 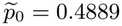 for (C). The first three columns **(A, B, C)** use the same nonlinear elastic model, with 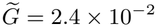 and *ϵ*_*c*_ = 0.04, whereas the last column **(D)** uses a linear elastic model, with 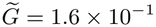 and *ϵ*_*c*_*→ ∞*. The corresponding pre-strain *ϵ*_0_ is indicated below 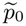 in each first-row plot. The first row **(A1, B1, C1, D1)** shows the contact area normalized by *β D*^2^ as a function of the compression amplitude *ξ*. The second row **(A2, B2, C2, D2)** shows the contact area normalized by *η D*^2^, where *η* is given by Eq. (32), with *φ*_0_ = 0.34, *k* = 0.37, and *τ* = 7.00 for (A, B, C), and *φ*_0_ = 0.25, *k* = 0.22, and *τ* = 7.00 for (D), consistent with the parameters provided in Table I. The dashed lines represent the general contact area model in Eq. (33), while the dash-dotted lines represent the linear approximation given by Eq. (34).

**FIG. A.7:**
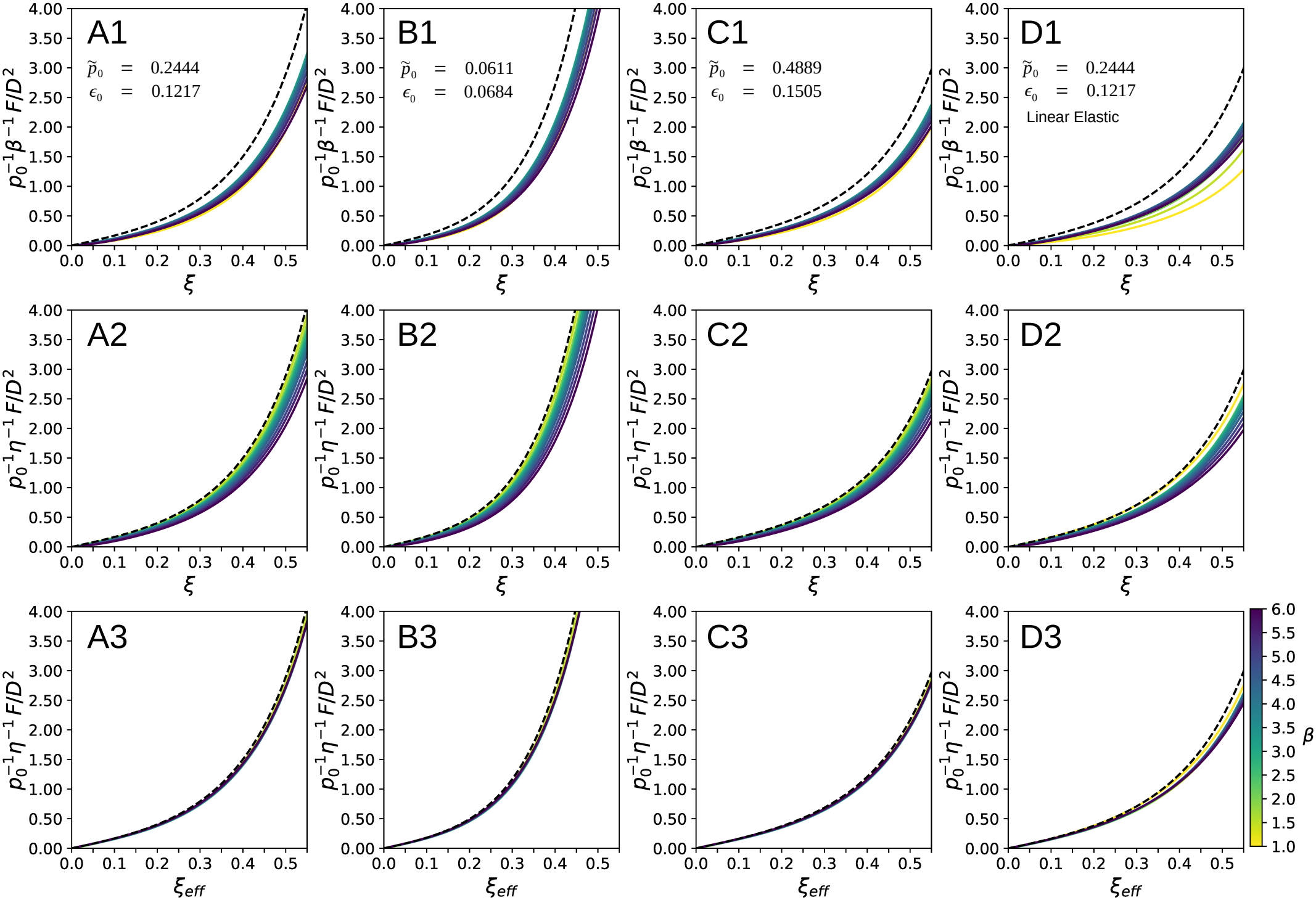
Renormalization of the loading force for large-bead compression. Data obtained from simulations using *Surface Evolver* with *R* = 34 *× D* and aspect ratios in the range 1 ≤ *β*≤ 6. The color of the solid lines indicates *β*, increasing from yellow to blue according to the color map (reverse *viridis*) used in Figs. 2B and 7 (color bar in the lower-right corner). The dashed line corresponds to the normalized force density 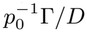 obtained for cylinders in the parallel-plate limit, Eq. (17); in this limit, the curves are unaffected by the renormalizations. Each column (A, B, C, D) corresponds to a different set of control parameters. The values of 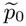 are indicated in the first-row plots: 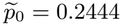 for (A, D), 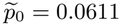 for (B), and 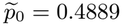 for (C). The first three columns **(A, B, C)** use the same nonlinear elastic model, with 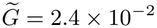 and *ϵ*_*c*_ = 0.04, whereas the last column **(D)** uses a linear elastic model, with 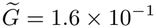and *ϵ*_*c*_*→ ∞*. The corresponding pre-strain *ϵ*_0_ is indicated below 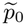 in each first-row plot. The first row **(A1, B1, C1, D1)** shows the force normalized by *p*_0_ *β D*^2^ as a function of the compression amplitude *ξ*. The second row **(A2, B2, C2, D2)** shows the force renormalized by *p*_0_ *η D*^2^, where *η* is given by Eq. (32), with *φ*_0_ = 0.34, *k* = 0.37, and *τ* = 7.00 for (A, B, C), and *φ*_0_ = 0.25, *k* = 0.22, and *τ* = 7.00 for (D), consistent with the parameters provided in Table I. The third row **(A3, B3, C3, D3)** shows the renormalized force as a function of the effective compression amplitude *ξ*_eff_.

**FIG. A.8:**
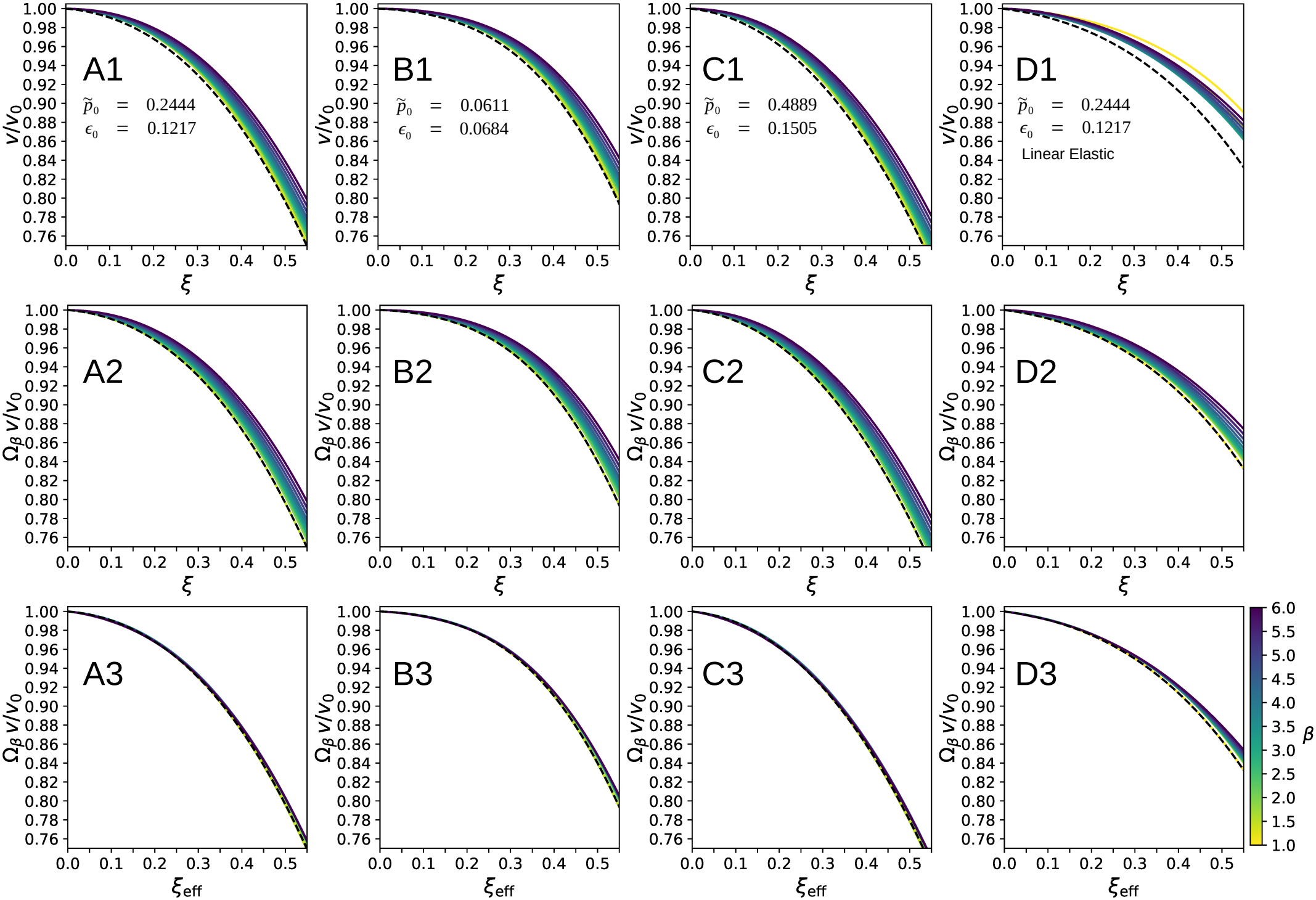
Renormalization of the volume for large-bead compression. Data obtained from simulations using *Surface Evolver* with *R* = 34 *× D* and aspect ratios in the range 1 ≤ *β* ≤ 6. The color of the solid lines indicates *β*, increasing from yellow to blue according to the color map (reverse *viridis*) used in Figs. 2B and 7 (color bar in the lower-right corner). The dashed line corresponds to the relative volume density *ρ/ρ*_0_, with 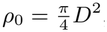, obtained for cylinders in the parallel-plate limit, Eq. (18); in this limit, the curves are unaffected by the renormalizations. Each column (A, B, C, D) corresponds to a different set of control parameters. The values of 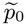 are indicated in the first-row plots: 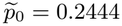 for (A, D), 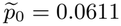 for (B), and 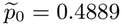 for (C). The first three columns **(A, B, C)** use the same nonlinear elastic model, with *G* = 2.4 *×* 10^−2^ and *ϵ*_*c*_ = 0.04, whereas the last column **(D)** uses a linear elastic model, with 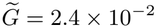 and *ϵ*_*c*_ *→ ∞*. The corresponding pre-strain *ϵ*_0_ is indicated below 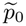 in each first-row plot. The first row **(A1, B1, C1, D1)** shows the relative volume *v/v*_0_ as a function of the compression amplitude *ξ*. The second row **(A2, B2, C2, D2)** shows the relative volume multiplied by the renormalization factor 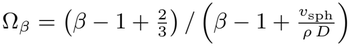. The third row **(A3, B3, C3, D3)** shows the renormalized relative volume as a function of the effective compression amplitude *ξ*_eff_.

**FIG. A.9:**
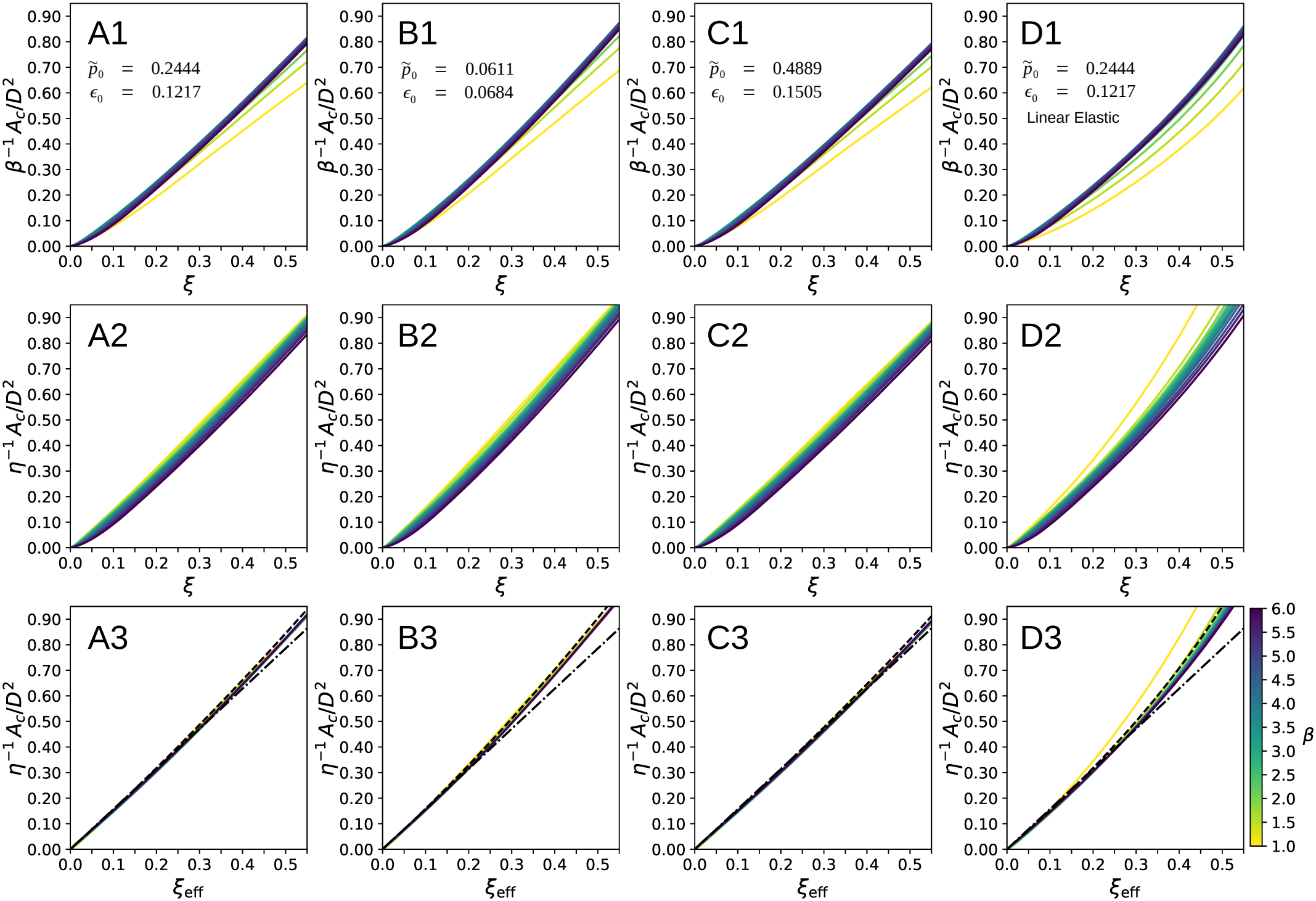
Contact area for large-bead compression. Data obtained from simulations using *Surface Evolver* with *R* = 34 *× D* and aspect ratios in the range 1 ≤ *β* ≤ 6. The color of the solid lines indicates *β*, increasing from yellow to blue according to the color map (reverse *viridis*) used in Figs. 2B and 7 (color bar in the lower-right corner). Each column (A, B, C, D) corresponds to a different set of control parameters. The values of 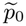 are indicated in the first-row plots: 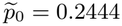 for (A, D), 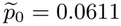 for (B), and 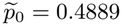 for (C). The first three columns **(A, B, C)** use the same nonlinear elastic model, with 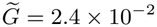 and *ϵ*_*c*_ = 0.04, whereas the last column **(D)** uses a linear elastic model, with 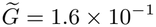 and *ϵ*_*c*_ *→ ∞*. The corresponding pre-strain *ϵ*_0_ is indicated below 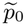 in each first-row plot. The first row **(A1, B1, C1, D1)** shows the contact area normalized by *β D*^2^ as a function of the compression amplitude *ξ*. The second row **(A2, B2, C2, D2)** shows the contact area renormalized by *η D*^2^, where *η* is given by Eq. (32), with *φ*_0_ = 0.34, *k* = 0.37, and *τ* = 7.00 for (A, B, C), and *φ*_0_ = 0.25, *k* = 0.22, and *τ* = 7.00 for (D), consistent with the parameters provided in Table I. The third row **(A3, B3, C3, D3)** shows the renormalized contact area as a function of the effective compression amplitude *ξ*_eff_. The dashed lines represent the general contact area model in Eq. (33), while the dash-dotted lines represent the linear approximation given by Eq. (34).

**FIG. A.10:**
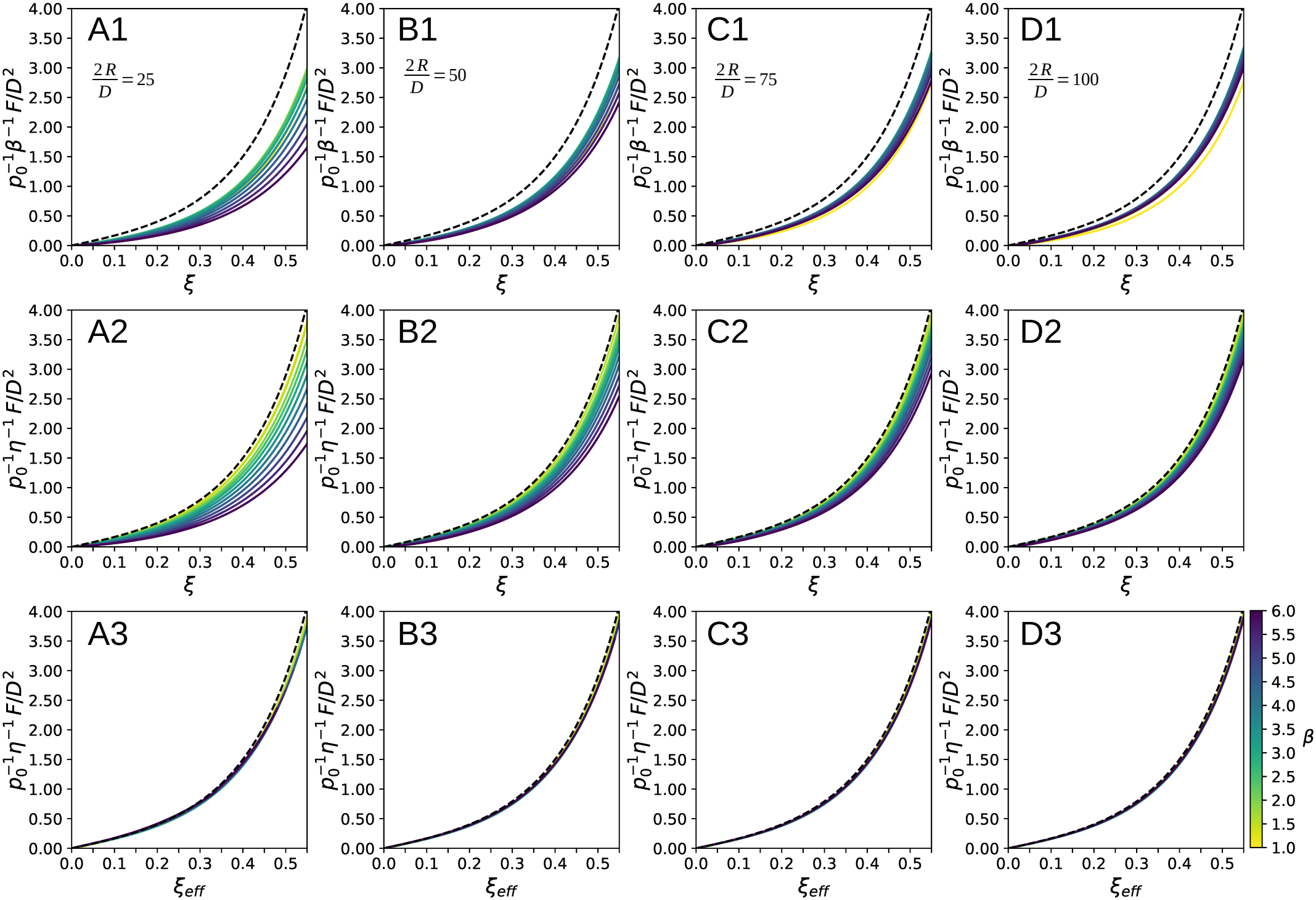
Renormalization of the loading force for different bead radii. Data obtained from simulations using *Surface Evolver* with control parameters 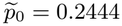 and *ϵ*_0_ = 0.1217, and a nonlinear elastic model (*ϵ*_*c*_ = 0.04). The data in each column (A, B, C, D) correspond to a different bead radius: *R/D* = 25*/*2 (panels A1, A2, and A3), *R/D* = 50*/*2 (panels B1, B2, and B3), *R/D* = 75*/*2 (panels C1, C2, and C3), and *R/D* = 100*/*2 (panels D1, D2, and D3). The solid-line colors indicate the capsule aspect ratio, 1 ≤ *β* ≤ 6, increasing from yellow to blue according to the reverse *viridis* color map used in Figs. 2B and 7 (color bar in the lower-right corner). The dashed line corresponds to the normalized force density 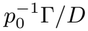 for cylinders in the parallel-plate limit, given by Eq. (17); in this limit, the curves are unaffected by the renormalizations. The first row (A1, B1, C1, D1) shows the force normalized by *p*_0_*βD*^2^ as a function of the compression amplitude *ξ*. The second row (A2, B2, C2, D2) shows the force renormalized by *p*_0_*ηD*^2^, where *η* is given by Eq. (32), with *φ*_0_ = 0.34, *k* = 0.37, and *τ* = 7.00, consistent with the parameters provided in Table I. The third row (A3, B3, C3, D3) shows the renormalized force as a function of the effective compression amplitude *ξ*_eff_.

**FIG. A.11:**
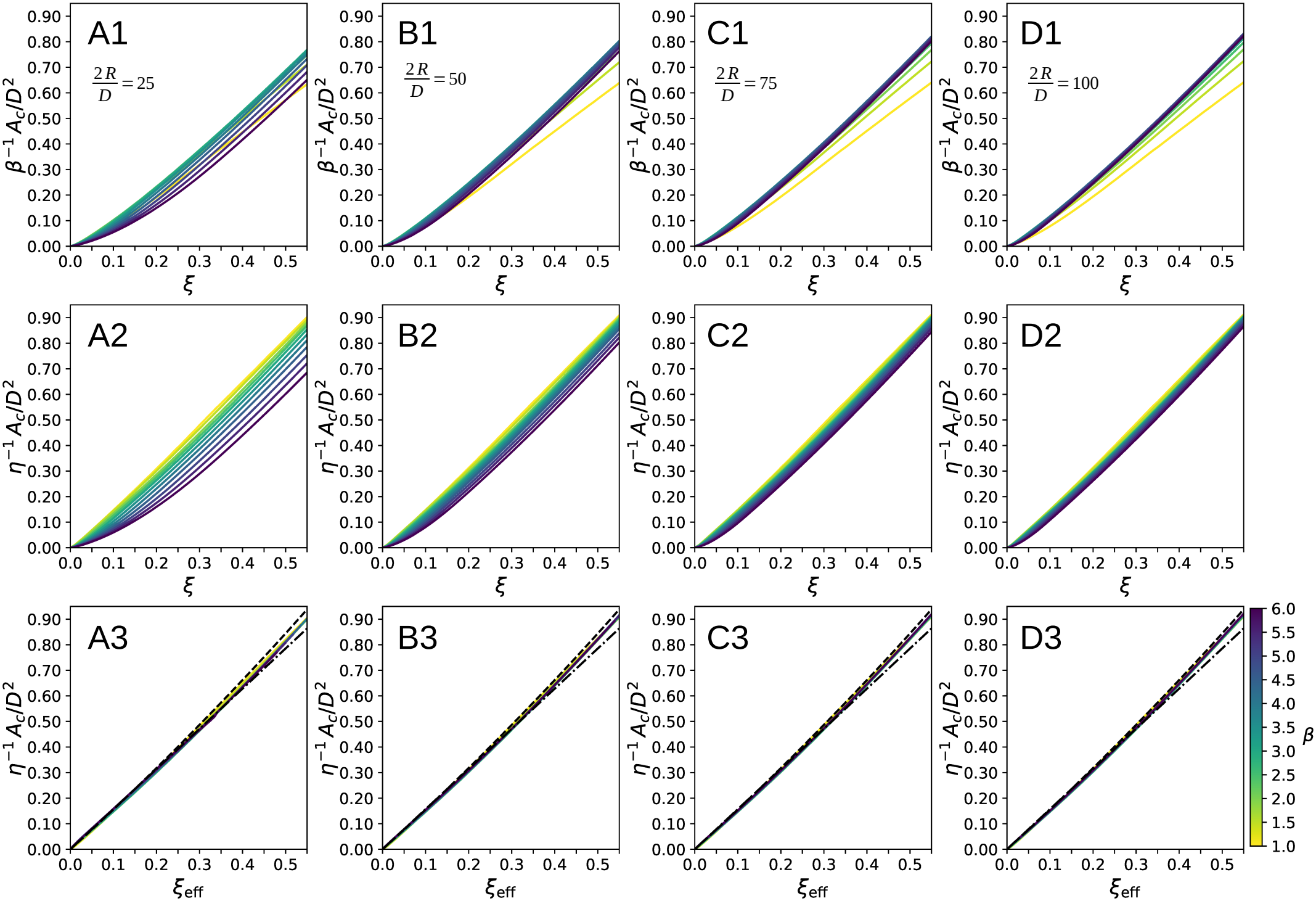
Contact area for different bead radii. Data obtained from simulations using *Surface Evolver* [5] with control parameters 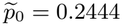 and *ϵ*_0_ = 0.1217, and a nonlinear elastic model (*ϵ*_*c*_ = 0.04). The data in each column (A, B, C, D) correspond to a different bead radius: *R/D* = 25*/*2 (panels A1, A2, and A3), *R/D* = 50*/*2 (panels B1, B2, and B3), *R/D* = 75*/*2 (panels C1, C2, and C3), and *R/D* = 100*/*2 (panels D1, D2, and D3). The solid-line colors indicate the capsule aspect ratio, 1 ≤ *β* ≤ 6, increasing from yellow to blue according to the reverse *viridis* color map used in Figs. 2B and 7 (color bar in the lower-right corner). The first row (A1, B1, C1, D1) shows the contact area normalized by *βD*^2^ as a function of the compression amplitude *ξ*. The second row (A2, B2, C2, D2) shows the contact area renormalized by *ηD*^2^, where *η* is given by Eq. (32), with *φ*_0_ = 0.34, *k* = 0.37, and *τ* = 7.00, consistent with the parameters provided in Table I. The third row (A3, B3, C3, D3) shows the renormalized contact area as a function of the effective compression amplitude *ξ*_eff_. The dashed lines represent the general contact area model in Eq. (33), while the dash-dotted lines represent the linear approximation given by Eq. (34).

